# Emergent digital bio-computation through spatial diffusion and engineered bacteria

**DOI:** 10.1101/2023.07.07.548081

**Authors:** Alex J.H. Fedorec, Neythen J. Treloar, Ke Yan Wen, Linda Dekker, Qing Hsuan Ong, Gabija Jurkeviciute, Enbo Lyu, Jack Rutter, Luca Rosa, Alexey Zaikin, Chris P. Barnes

**Author notes:** These authors contributed equally to this work (Alex J.H. Fedorec, Neythen J. Treloar, Ke Yan Wen).

## Abstract

Building computationally capable biological systems has long been an aim of synthetic biology. The potential utility of biocomputing devices ranges from biosafety and environmental applications to diagnosis and personalised medicine. Here we present work for the design of bacterial computers which use spatial patterning to process information. Our computers are composed of bacterial colonies which, inspired by patterning in embryo development, receive information in the form of diffusible morphogen-like signals. A computation is encoded by the physical locations of the input sources of morphogen and the output receiver colonies. We demonstrate, mathematically and with engineered *Escherichia coli*, the simple digital logic capability of single bacterial colonies and show how additional colonies are required to build complex functions. Inspired by electronic design automation, an algorithm for designing optimal spatial circuits computing two-level digital functions is presented. This enhances the capability of our system to complex digital functions without increasing the biological complexity. We extend our experimental system to incorporate biosensing colonies as morphogen sources, demonstrating how a diagnostic device might be constructed. Our approach will open up new ways to perform biological computation, with applications in bioengineering, biomaterials and biosensing. Ultimately, these computational bacterial communities will help us explore information processing in natural biological systems.

## Introduction

From the demonstration of DNA computing in the early 1990s, people have wondered how biological substrates can be used for computation. Engineering biological systems capable of computation has also long been a goal of synthetic biology. The first synthetic biology papers engineered a toggle switch [1], oscillator [2] and autoregulation [3], which can be used as fundamental components in engineering a computer [4]: memory, clock and noise filter. The most common paradigm in biological computation to date has been the engineering of programmable logic into gene regulatory networks (GRNs) inside cells [5]. The current state-of-the-art are automated design tools such as Cello [6]–[8], which compiles a digital function – written in a Verilog-like language – into its corresponding GRN. This enables a comprehensive suite of three input digital functions to be constructed.

However, for each new function, a new genetic circuit must be built which means extensive genetic engineering. Furthermore, there are practical limitations on the complexity of circuits that can be programmed into a single cell due to a limited number of available, orthogonal components, and metabolic burden. Because components in a cell cannot be directly wired together, we cannot reuse gene regulators without interference, which limits the Cello system to 12 regulators [6]. In contrast, metabolic burden becomes a problem when the resource requirements of a GRN in a cell approaches the capacity of the cell’s metabolism. Under high metabolic load a cell may exhibit increased mutation rates due to stress responses [9] and a competitive advantage to losing or mutating the programmed circuit [10]. These factors reduce the stability of a programmed GRN and place a practical upper limit on the complexity of the constructed function.

To overcome these limitations, computational communities of microbes have been developed in liquid culture. This allows the division of a complex function into smaller, isolated modules which can be programmed into different populations and integrated using inter-population communication. Two-and three-input logic gates have been implemented in yeast liquid culture [11], and microbial consortia capable of four-input digital logic [12] have also been built in this way. Algorithms for the optimal design of digital functions in liquid communities have been developed [13]. However, as with the orthogonality requirements of GRNs within single cells, building microbial communities that communicate in liquid culture requires the use of a unique communication molecule for each “wire” and the avoidance of crosstalk between molecules. This has previously been termed the “wiring problem” of wet computational circuits [14].

The limitations associated with monoculture computing and distributed communities in liquid culture can be overcome by producing logic gates based on spatially arranged, communicating bacterial colonies in solid culture. The GRNs inside each colony can be simple and isolated, naturally overcoming the limitations of burden and non-orthogonality. Furthermore, the signal produced by a colony is spatially localised, meaning that other colonies can be insulated from it by placing them outside of the signal range, largely avoiding the wiring problem associated with liquid culture communities. Early work using colonies on solid media combined bacterial logic gates with spatial signalling by using four different quorum molecules to connect bacterial NOR gates [15]. Another work decomposed digital functions into the sum of spatially separated modules, where each module is a negated sum of IDENTITY and NOT functions of the inputs [16]. Subsequent research used the spatial arrangement of sender cells and modulator cells where a function is printed in a branched architecture on a piece of paper and was used to implement functions such as a three-input parity bit [17]. That work enabled the printing of digital functions using stamps and “cellular ink”, and demonstrated that the printed functions were storable for later use.

In this work, we combine a small set of simple inducer activation functions [18], [19] with spatial structure as a control parameter, to produce designs for modular, easily programmable bacterial computers. In contrast to previous work, much of the information processing is encoded in the spatial patterning of colonies. The use of programmable spatial patterns and the range of patterning functions is unique to our approach and enables us to program arbitrary digital functions with minimal genetic engineering. By using the distance between colonies as a design parameter, we can use each activation function as a flexible module that encodes different logic depending on its position relative to its inputs. Unlike designing and building a complex GRN into a cell, a new pattern can easily be produced using automated pipetting robots.

Our computational devices consist of bacterial receiver cells positioned on a plate relative to sources of diffusible molecules. Each source represents an input to the computation and the receivers produce a fluorescent output that represents the result of the computation. Because communication by a diffusible molecule is dependent on distance, the bacterial computer can be programmed by specifying the spatial pattern of the receivers and the sources. First, we demonstrate the position dependence of our circuits using a lawn of receiver cells and droplets of IPTG as diffusible inputs. Then we demonstrate the construction of some two-input digital functions using colonies of receiver cells. We show that each receiver can be programmed to encode many different two-input logic gates by changing their position relative to the inputs. We present a mathematical framework to understand these results and make mathematical statements of its capability. We then develop an algorithm to optimise the programming of complex digital functions distributed over multiple receivers. A key feature of this algorithm is that it encodes a given digital function in a form that can be implemented in one level of biological signalling, which minimises the complexity of the spatial functions and greatly reduces the required genetic engineering. This approach also guarantees that any function can be implemented using only one communication molecule, minimising the wiring requirements. Finally, we demonstrate the use of biosensor colonies as sources of the diffusible molecule, enabling computation based on multiple environmental inputs, providing a simple proof-of-concept for future applications such as medical diagnostics or pollution testing.

## Results

### Analogue-to-digital conversion of interacting diffusion patterns allows for spatial computing

Inputs and outputs of a digital function can be ON (1) or OFF (0). For a two-input digital function this results in four possible input states - 00, 01, 10, 11 - each of which can produce a 0 or 1 output. There are sixteen unique ways of mapping the four input states to the binary output, resulting in a total of sixteen unique two-input logic gates. In our computation paradigm, inputs are sources of diffusible molecules, with 0 and 1 indicating the absence or presence of each source respectively. These inputs produce a unique diffusion field for each of the four input states. Bacterial “receiver” cells, capable of sensing and responding to the concentration of the diffusion field at their specific locations, map the input states to binary outputs.

Here, we consider four ways in which receiver cells can respond to increasing concentrations of diffusible molecule: a “highpass”, where the cells are switched on above a certain concentration; a “bandpass”, where cells are only on at intermediate concentrations; and their inverses, a “lowpass” and “bandstop”. In a homogenous field of receiver cells, the diffusion field produced by the two inputs will trigger a response in each of the cells depending on where they are located, resulting in a specific response pattern. Using a toy model simulated with the finite difference method [20], we demonstrate that a diffusion field, interpreted using our set of activation functions, produces digital logic (Figure 1A, B). For each of the input configurations, a simulation was performed in which each input is a source of diffusible molecule added at the beginning of the simulation. The four activation functions were applied to the resultant diffusion fields and the digital functions produced at each position were determined. This results in a map where the colour of each region represents the logic gate that would be encoded if a receiver cell was placed in that region. These simulations show that with the bandpass and bandstop alone, it is possible to encode all sixteen two-input digital logic gates. It is also clear that the logic gate encoded by a given receiver is dependent on its position relative to the inputs.

**Figure 1:**
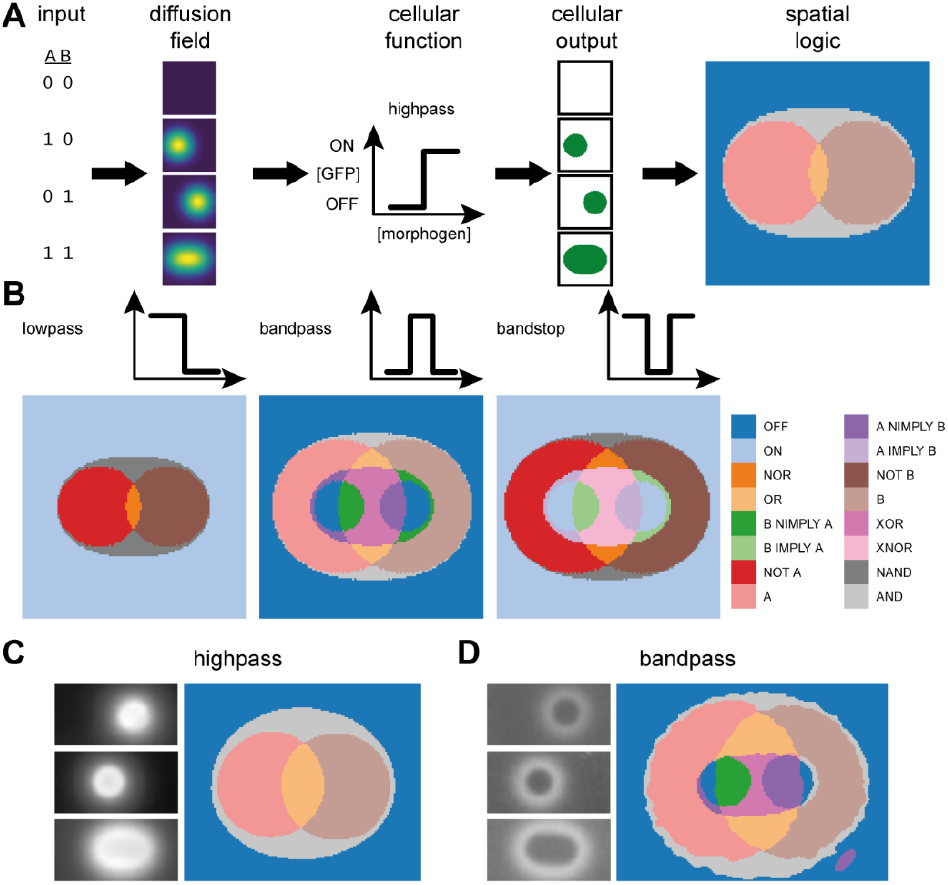
Logic gates can be produced from a combination of spatial position and engineered response function. A) Given two possible digital inputs, four combinations can be produced. If each input corresponds to a diffusible molecule, each input configuration produces a unique gradient pattern. Cells are engineered to respond to the concentration, [I], of the diffusible molecule in a specific way – in this example, turning on above a threshold. Cells at each location in the environment turn on or off depending on the input configuration and their engineered function. Each location can then be mapped to a specific logic gate given the cells’ responses. B) Different response function will produce different logic gates. All sixteen two-input logic gates can be produced with bandpass and bandstop response functions. C & D) Response of a lawn of highpass and bandpass receiver cells induced with droplets of IPTG at either A, B, or A and B input locations. Images are of plates after 20 hours. The images are processed and a digital function is assigned to each pixel.

### Engineered *E. coli* can generate predicted digital logic from diffusion fields

To implement the system experimentally, we utilised genetic circuits which respond to IPTG to form either highpass or bandpass receiver responses through the expression of GFP [19]. Their dose responses were characterised in liquid culture (Supplementary Figure 3). We sought to understand how the IPTG diffuses and how the bacteria growing on an agar substrate respond. Agar plates were seeded with our engineered cells and we dispensed droplets of IPTG at different concentrations (Supplementary Figure 4A). The diffusion field created by the IPTG droplet depends on its concentration and the time allowed for it to diffuse. One might, therefore, expect the diameter of the ring formed by the bandpass cells to change with concentration and time. Indeed, with higher inducer droplet concentrations, the ring size increases. However, the observable position of the ring is temporally static, indicating that at some point – perhaps upon reaching a particular growth phase – the cellular response is established and fixed.

With an understanding of the diffusion field and the response of our cells, we can design spatial patterns of inputs which can produce specific logic outputs. Indeed, with this information we can place droplets of IPTG inducer at defined positions on a field of cells and demonstrate that all the predicted regions present for the highpass and bandpass receiver can be observed in practice (Figure 1C, D). Simply by changing the distance between the two inputs, we change the position of specific logical responses to those inputs (Supplementary Figure 4B).

### Precisely positioned engineered bacterial colonies serve as digital function outputs

Given that we can observe different digital functions at specific positions within a field of cells, we hypothesised that it should also be possible to place colonies at defined locations and interpret their output response as a digital logic gate. The use of colonies instead of a field of cells simplifies the read-out, as the position from which to read the output has been pre-defined by the colony placement. Further, it enables the use of different receiver cell types at different positions which will allow us to construct more complex logic gates.

We used a liquid handling robot to dispense droplets of culture that form into colonies (Figure 2A). This restricts us to positions defined by a 384-well microtitre plate layout. We characterised the responses of bandpass and highpass colonies at defined distances from droplets of different concentrations of IPTG input (Supplementary Figure 3). With this data we were able to fit a reaction-diffusion model which incorporates growth of the bacterial colonies, diffusion of the IPTG inputs, and the dynamics of the transcriptional networks within the cells (Supplementary Information 4 and Supplementary Figure 12).

**Figure 2:**
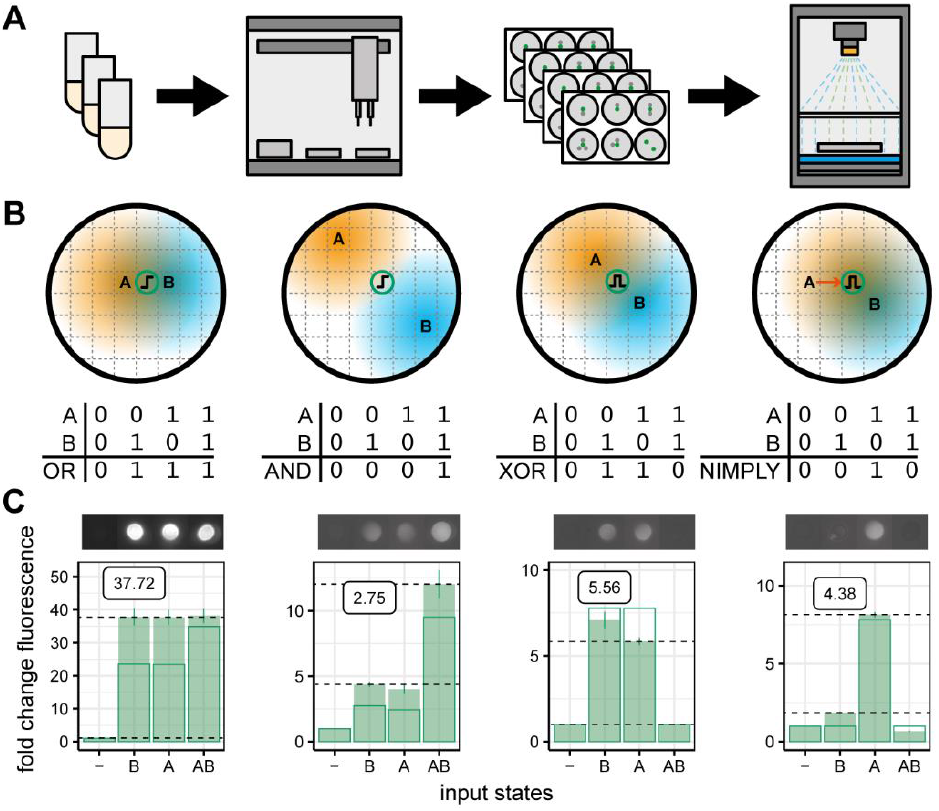
Biological implementation of the spatial computing framework. A) Receiver colonies and IPTG inputs were dispensed at pre-defined positions using a liquid handling robot, and imaged for growth and fluorescence. B) Predicted spatial layouts that will give rise to OR, AND, XOR and NIMPLY logic responses. Letters indicate the position of the respective IPTG inputs, circles containing highpass or bandpass cartoon indicate location of output colonies. Coloured regions represent the diffusion of the IPTG from the input locations. C) Colony responses to all input configurations at 20 hours. Images show a representative colony for each configuration (highpass OR exposure=40000, intensity=3, all other gates exposure=60000, intensity=5). Solid bars show the mean fold change in fluorescence relative to a colony with no IPTG inputs. Error bars show the standard error of the mean for three replicates. Outlined bars show model predications. The gate score displayed on the plot shows the least fluorescent ON state divided by the most fluorescent OFF state (shown with dashed horizontal lines).

We designed spatial patterns of inputs and receiver colonies that are capable of producing the four, non-trivial, two-input logic gates achievable with the highpass and bandpass (Figure 2B). We then implemented these spatial patterns experimentally in 6-well plates using droplets of IPTG for inputs and colonies of bacterial receivers as outputs (Figure 2C). The growth and fluorescence of each colony was determined using a custom imaging system (Methods). The fold change in fluorescence was calculated and each gate was scored by dividing the lowest ON state by the highest OFF state [6]. All of the gates perform their predicted function, though there are deviations from the model predicted fluorescence levels. One can clearly see the high fluorescence output of the highpass colonies compared with the bandpass colonies, leading to a very high score for the highpass OR gate. The AND gate does not perform as well and the reason for this can be intuited from the dose response curve of the highpass (Supplementary Figure 3). A doubling of inducer concentration – as when moving from one input being present to both inputs present – cannot result in a shift from minimal expression to maximal expression due to the shallowness of the dose response curve. The best we can achieve with our highpass is approximately 3-fold fluorescence change.

### An abstract representation of receiver response to signalling gradients facilitates an analysis of realisable digital functions

Up to this point, the logic gates that we have implemented have been simple enough that sensible positions for inputs and outputs can be inferred from the receiver characterisation data. However, as we wish to construct more complex logical functions, a computational approach is required. To this end, we developed an abstract representation of the inducer concentration seen by a receiver for each input state, allowing us to investigate the capabilities of the system (Supplementary Information 1.1).

We simplify the description of the spatial arrangement of the receiver and sources by considering a single dimension; the signal concentration observed by the receiver. The presence of a source of inducer increases the signal concentration seen by the receiver by an amount proportional to the distance between the receiver and source. By moving the receiver closer to a source, we increase the concentration seen by the receiver. To reflect this change, the input states that include that source move further up our signal concentration dimension. Similarly, moving away from a source does the inverse. Figure 3A shows that moving a receiver relative to two sources changes the order of the input states on the signal concentration dimension.

**Figure 3:**
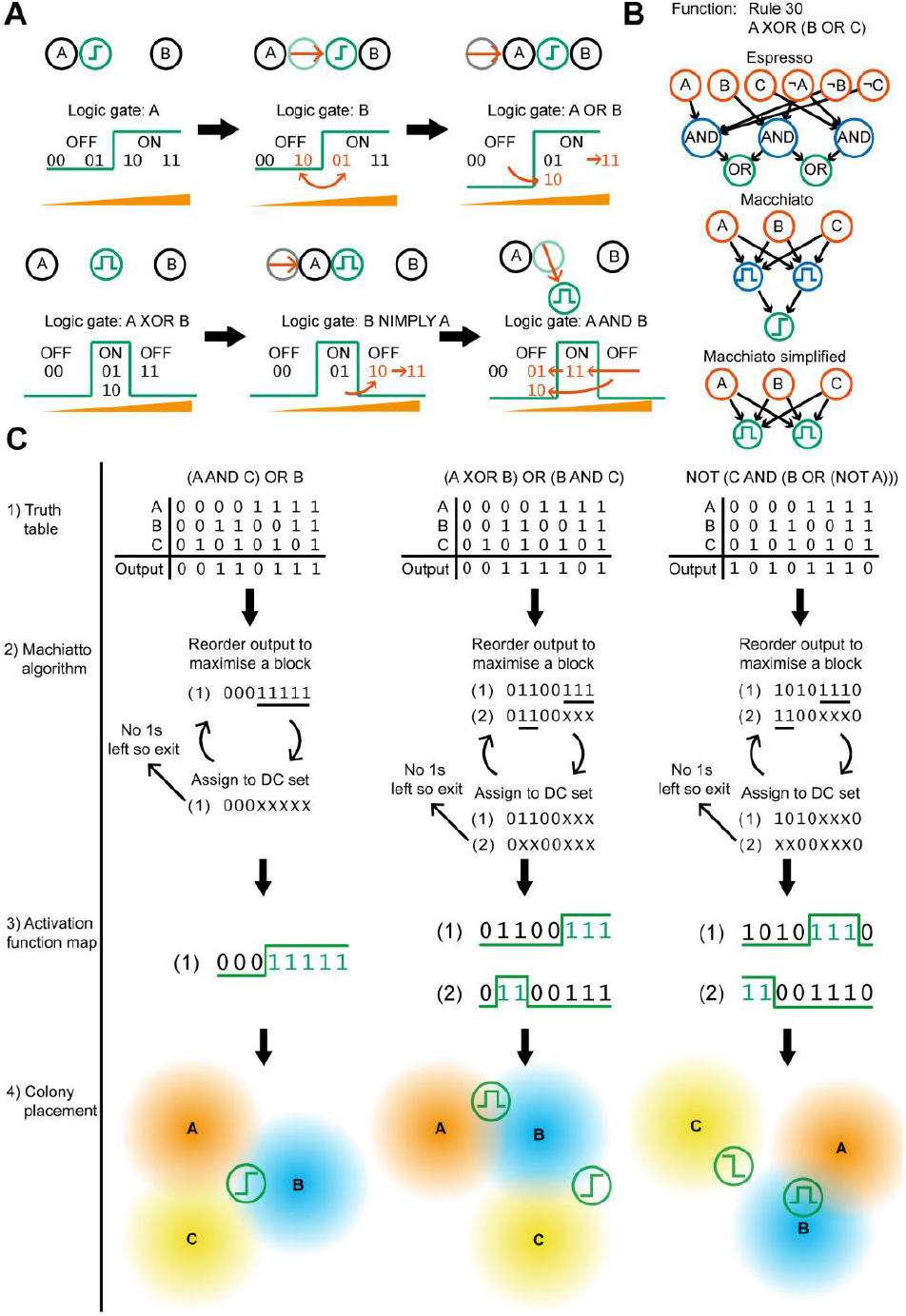
Algorithmic determination of input and output patterning. A) Demonstration of how varying positions and activation functions results in programmable digital logic and how this is represented in our framework. B) A digital logic function simplified by the Espresso algorithm (top), Macchiato algorithm with two levels of biological signalling (middle) and Macchiato with one level of biological signalling (bottom). C) The stages of the Macchiato algorithm on three examples, from left to right; 1) a truth table is supplied as input, 2) the input states are rearranged to minimise the number of blocks while obeying the constraints imposed by the system (DC = “don’t care”), 3) activation functions are mapped onto the blocks, 4) spatial configuration of nodes are produced that results in the truth table, if any of the green output colonies are activated the output of the function is ON.

We represent the activation functions – highpass, lowpass, bandpass and bandstop - as boundaries partitioning these ordered input states into ON and OFF boxes (Supplementary Information 1.2). Figure 3A (and Supplementary Figure 1) shows how this partitioning of input states results in logic output. By moving the receiver and sources, we move the input states in and out of the ON and OFF boxes. In the way we can change the logic gate encoded by different patterns of inputs and receiver. Using this representation, we can enumerate of all the distinct logic gates that are possible, given a number of inputs and the available receivers, while obeying physical constraints. All two-input digital functions can be computed spatially using a single receiver, provided we have receivers with response functions given in Figures 1A and B (Supplementary Information 1.3), corroborating the simulation results.

### Two level logic enables the construction of arbitrary digital functions

We extended our theoretical analysis of spatial computing to three-input logic gates and found that we can only encode 152 of the 256 possible three-input logic gates with a single receiver output (Table 1, Supplementary Table 2). This demonstrates a single receiver output is not sufficient to enable the construction of arbitrarily complex digital functions. To overcome this limitation, we developed an algorithm for the optimisation of two-level spatial digital logic. In electronic design, logic minimisation is used to simplify a digital circuit into a minimal canonical form. The Espresso algorithm is a heuristic optimiser that attempts to minimise a function into the minimal set of OR of ANDs, where each AND term can contain any of the inputs or their negations [21] (Figure 3B, top).

**Table 1:** Receiver activation function requirements for three-input digital function computation. The second column shows the number of functions not computable. The last two columns gives the number of functions with 1 and 2 required receivers. (HP: highpass, LP: lowpass, BP: bandpass, BS: bandstop).

| Activation functions | Not possible | Number of receivers |  |
| --- | --- | --- | --- |
|  |  | 1 | 2 |
| BP, BS, LP, HP | 0 | 152 | 104 |
| BP, BS, LP | 0 | 152 | 104 |
| BP, BS, HP | 0 | 152 | 104 |
| BP, LP, HP | 0 | 95 | 161 |
| BS, LP, HP | 155 | 95 | 6 |
| BP, BS | 0 | 152 | 104 |
| BP, LP | 0 | 95 | 161 |
| BP, HP | 128 | 76 | 52 |
| BS, LP | 174 | 76 | 6 |
| BS, HP | 155 | 95 | 6 |
| LP, HP | 155 | 38 | 63 |
| BP | 128 | 76 | 52 |
| BS | 174 | 76 | 6 |
| LP | 237 | 19 | 0 |
| HP | 237 | 19 | 0 |

Taking inspiration from the Espresso algorithm we have developed a novel algorithm, named the Macchiato algorithm, which finds the minimal set of receivers that encode a given digital function. The Macchiato algorithm takes the truth table of a multi-input, one-output function and returns the approximate minimal set of bandpass, highpass, lowpass and bandstop receivers that are required to construct the digital function and the required mapping of input states for each receiver. The minimal set of receivers constitutes the analogue of the AND layer and the mapping represents the connectivity to the inputs. The OR can then be performed using a highpass receiver that activates if any of the previous layer is activated (Figure 3B, middle) or it can be done implicitly in the sense that if any individual receiver is ON we take the output as ON (Figure 3B, bottom). It is for this reason we focus on an analogue of the OR of ANDs (sum of products) form of a digital function, as we gain the flexibility to encode two-level digital functions using only one level of biological signalling.

For each three-input gate the number of bacterial colonies required to build the gate was calculated for three different sets of available activation functions (Table 1, Supplementary Table 3). With all the activation functions, all three-input logic gates are possible and only require one or two colonies. If we remove the bandstop from the available activation functions we retain the capability to do all the three-input logic gates, however a larger proportion now require two bacterial colonies. If we also remove the bandpass activation we lose the capability to compute the majority of the three input logic gates.

The final step is to determine the optimal spatial positions of the inputs and receivers relative to one another. To do this, we use a calibrated finite difference model and an evolutionary algorithm, optimising the difference in expected fluorescence between ON and OFF states (Figure 3C).

### Designing three-input spatial logic gates using the Macchiato algorithm

We demonstrate our approach by constructing a selection of three-input logic gates (Figure 4). These are designed using the Macchiato algorithm to find the minimal set of highpass and bandpass receivers that will encode the given function. We chose logic gates that cover a range of complexity if one were to implement them using state-of-the-art gene regulatory network designs [6]: 0×7F = 27 genetic parts, 0×37 = 32, Multiplexer = 38, Rule 30 = 44, Majority = 49. All of the logic gates function as expected. However, as with the two-input gates demonstrated above, the logic gates that require AND functionality from the highpass receivers – for example 0×37 requires A AND C behaviour - score lower than the other logic gates.

**Figure 4:**
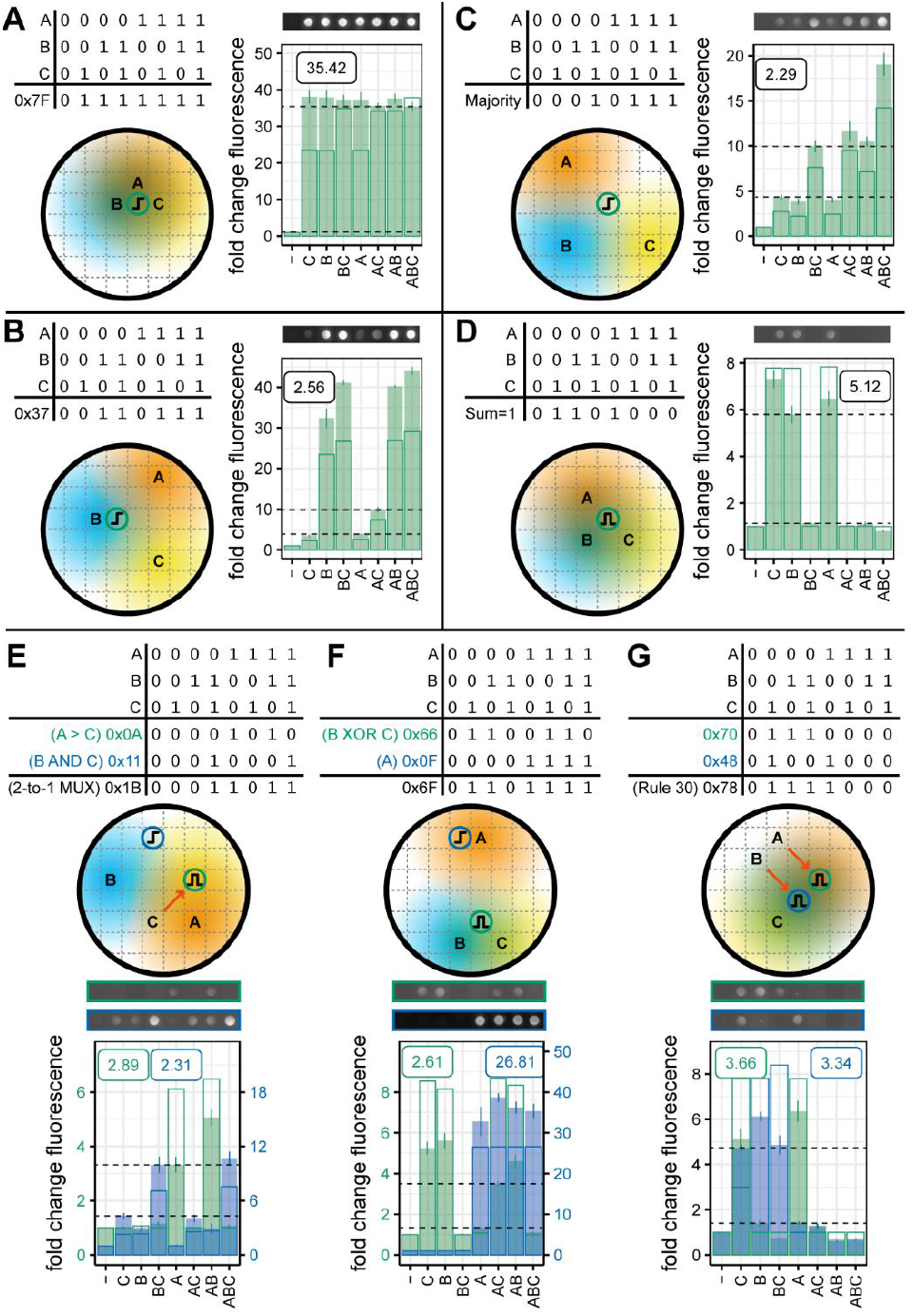
Computing three-input spatial logic. A selection of three-input logic gates with their output. Predicted spatial layouts that will give rise to 0×7F, 0×37, Majority, and sum = 1 functions. Circles containing letters indicate the location of IPTG inputs and the shaded regions represent the diffusion of the IPTG. Circles containing highpass or bandpass cartoon indicate location of output colonies. Colony images show representative fluorescence intensity at 20 hours (0×7F and 0×37 exposure=40000, intensity=3, Majority and Sum=1 exposure=60000, intensity=5). Solid bars show the mean fold change in fluorescence relative to a colony with no IPTG inputs. Error bars show the standard error of the mean for three replicates. Outline bars show model predications. The gate score displayed on the plot shows the least fluorescent ON state divided by the most fluorescent OFF state (shown with dashed horizontal lines).

We selected three logic-gates that require two output colonies, as it is not possible to minimise the output from the truth table to a single block. For example, in Figure 4E, a bandpass receiver produces A > C (0×0A) and a highpass receiver produces B AND C (0×11); the OR of these two outputs produces IF C THEN B ELSE A, otherwise known as a 2-to-1 multiplexer with C as the selector (0×1B). The digital function, again, performs as expected, however, the scores for both receiver colony functions are quite low. The highpass is performing an AND operation which does not score highly, as previously stated. In addition, the A input position relative to the bandpass receiver is not in its ideal position, resulting in slightly lower activation. This positioning was probably produced by our algorithm to increase the distance of A from the highpass receiver, to prevent unwanted activation. Similar considerations likely result in the lower performance of the B XOR C operation in the 0×6F gate (Figure 4F).

### Using biosensors as inputs for spatial digital logic

Up to this point, we have used robotically placed droplets of diffusible inducer as our function inputs. This is a rather restricted use case, limiting us to computations with a single input molecule and requiring printing of the computational device immediately prior to performing the computation. In order to remove these limitations, we move to a paradigm in which biosensing colonies act as sources of the diffusible molecule; producing it when triggered by the presence of an environmental input.

We constructed two biosensing *E. coli* strains that produce the diffusible quorum sensing molecule 3OC6-HSL, in response to chemical stimuli: arabinose and lactate. In addition, we built new receiver *E. coli* strains that respond to 3OC6-HSL with highpass and lowpass activation functions. The senders and receivers were characterised on solid culture (Supplementary Figure 6, 7). A lowpass and highpass receiver were selected that responded at similar inducer concentrations.

From the characterisation data, we determined patterns of sender and receiver colonies that produce the four non-trivial two-input logic gates that are achievable with highpass and lowpass outputs (Figure 5). Two of these – NAND and IMPLY – require two output colonies, which are computed from combinations of NOT and IDENTITY functions using the OR paradigm described previously (Figure 5D, E). A lower maximal fold-change for the highpass receiver, limits the score that can be achieved for gates using this strain as an output, however, all of these gates are successfully demonstrated.

**Figure 5:**
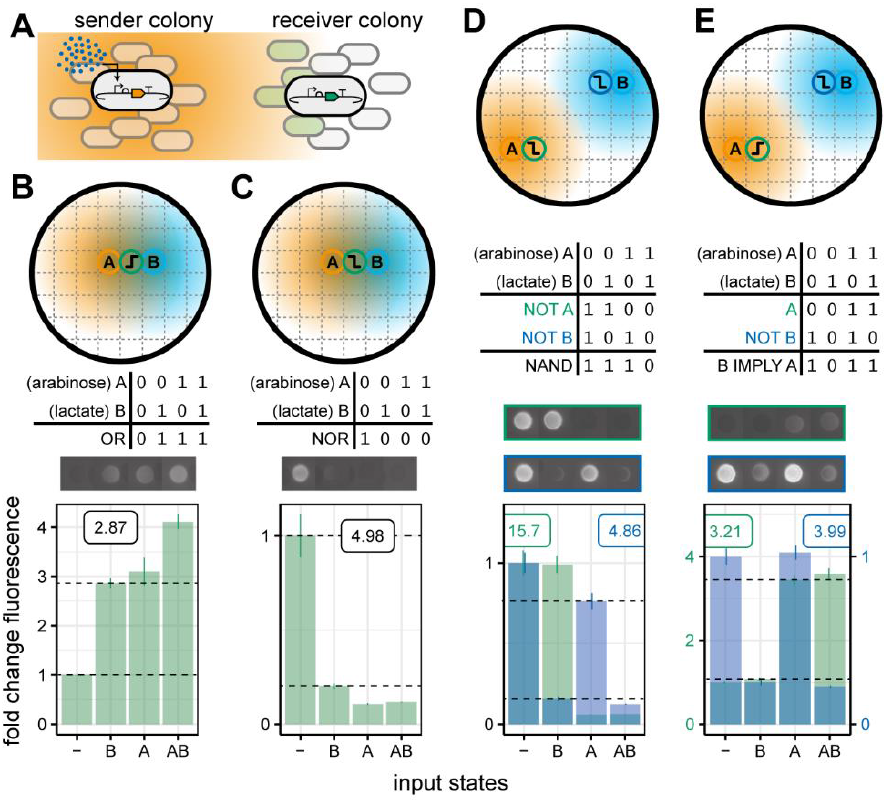
Biosensor colonies as morphogen sources for spatial computation. A) Biosensors, engineered to produce 3OC6-HSL in response to an environmental stimulus, were used as digital 5 inputs. B-E) Predicted spatial layouts that will give rise to OR, NOR, NAND and IMPLY logic responses. Circles containing letters indicate the position of the biosensor input colonies, circles containing highpass or lowpass cartoon indicate location of output colonies. Coloured regions represent the diffusion of 3OC6-HSL from the input colonies. Colony images show representative fluorescence intensity at 20 hours (all gates exposure=60000, intensity=5). Solid bars show the 10 mean fold change in fluorescence relative to a colony with no inducer inputs to the senders. Error bars show the standard error of the mean for three replicates. The gate score displayed on the plot shows the least fluorescent ON state divided by the most fluorescent OFF state (shown with dashed horizontal lines).

## Discussion

We have shown that interacting spatial morphogen gradients, combined with analogue-to-digital conversion, can encode complex biological computations. In our system, digital logic functions were built by patterning bacterial colonies, engineered to respond to diffusible molecules. We have proven that a single computing colony is capable of being programmed with any two-input logic gate by altering its position relative to its inputs or replacing it with an alternative activation function. This was demonstrated using both lawns of bacteria and precisely positioned bacterial colonies. The transition from lawns to colonies is important as it simplifies the measurement of the output; it is easy to measure the fluorescence of a colony but knowing which region of a lawn encodes a function is non-trivial. Further, the use of colonies allows us to use bacteria with different activation functions in different positions, enabling the construction of multi-output and multi-layered computations.

Additional complexity is necessary for the implementation of all three-input logic gates. We proposed an extension to our method, inspired by two-level electronic digital logic optimisation. This required us to develop an algorithm for an analogous spatial digital optimisation procedure to find the minimal set of bacterial colonies required for a given function. We extensively analysed this technique to find the requirements on the activation functions and provide a breakdown of the computational capabilities of the approach.

Incorporating sender colonies, capable of producing the diffusible molecule that the receivers respond to, expand the applicability of our computing system. Using biosensing senders, that are triggered by environmental inputs, enables the development of computational devices that can be used for medical or environmental diagnostics. The use of senders removes another limitation; directly printing the diffusible molecule inputs, as we did in Figures 1, 2 & 4 with IPTG, starts the computation immediately as the molecule begins to diffuse. By printing with sender colonies at the locations of the input sources, we can pre-prepare devices and the computation will only begin once the environmental stimuli is added. There has been great interest over the last decade in pre-prepared, bio-engineered, diagnostic devices [22].

In this work we have taken advantage of genetic circuits that have already been engineered to produce two of our required activation functions [23]. However, in order to generate all possible digital functions we would require a bandstop or lowpass genetic circuit. Further, the genetic circuits that we do have could be further optimised to produce a more digital response to stimuli [24]. This would provide cleaner transitions between ON and OFF regions, improving the signal-to-noise of our additive functions. Although our approach theoretically enables us to produce all three-input logic, as we increase the number of input and output positions, the connectedness is likely to make some gates unfeasible in two dimensions i.e. moving one input relative to one receiver also moves it relative to all other receivers. Moving to three-dimensional devices would relieve this limitation, and provide an avenue for the development of computational biomaterials [25], [26].

In comparison with previous, related approaches [15], [16] our system allows the construction of more complex functions with less biological complexity. This is because our choice of activation functions drastically offloads complexity from the internal cell dynamics to the morphogen field. This also enable a modular design that requires no genetic engineering to create a new biological computer once the initial set of activation functions have been produced. Even more compellingly, our approach is simpler than a state-of-the-art method to produce these functions in a gene regulatory network [6]. These properties make our platform an appealing approach for biological computing and potential real-world applications.

## Author contributions

Study conception and design: AJHF, NJT, KYW, CPB

Funding and supervision: AZ, CPB

Manuscript writing: AJHF, NJT, KYW, CPB

Theoretical results: AJHF, NJT, EL

Mathematical modelling: AJHF, NJT, LR

Experimental work: AJHF, KYW, LD, QO, GJ, JR, LR

