## Supplementary Information for "Emergent digital bio-computation through spatial diffusion and engineered bacteria"

### Supplementary Information: Emergent digital biocomputation through spatial diffusion and engineered bacteria

### 1 Theoretical foundations

#### 1.1 Representation of input states on a single ‘concentration’ dimension

Using two-input logic as an example, we consider the four possible input states – 00, 01, 10, 11 – as each lying on a line of increasing signal concentration seen by the receiver. The 00 state will always be the lowest concentration as we cannot reduce the concentration by adding an input. Similarly, the 11 state will always be the highest concentration seen by the receiver. However, the ordering of the 01 and 10 states will depend on the distance between the receiver and each of the inputs; when the are equidistant, 01 and 10 produce the same concentration seen by the receiver. However, if a receiver is one space away from input A and two spaces away from B, 10 will have a higher concentration than 01 and the order will be as above. If instead the receiver is next to B rather than A, the concentration seen by the receiver when B is on is higher than when A is on, and so the order will switch to 00, 10, 01, 11. The relative concentrations of the input states, and therefore their order, can change depending on the relative position of the receiver and the inputs (Figure 3A).

There are some constraints on the relative concentrations of signalling molecules which must hold for all possible receiver positions. For two inputs these are:

$$I_{00} \leq I_{01} \leq I_{11}$$
$$I_{00} \leq I_{10} \leq I_{11}$$
$$I_{ab} \geq 0 \; \forall \; a, b \in \{0, 1\}$$

where $I_{ab}$ is the concentration of signalling molecule at the receiver for input state $ab$. This imposes constraints on the allowable orderings of the input states.

#### 1.2 Activation functions partition the state space

With input states placed on a dimension of increasing signal concentration, we can conceptualise the activation functions engineered into the receiver cells as partitioning the state space into ON and OFF regions. For example, the highpass function remains in an OFF state at signal concentrations below a threshold, and switches to an ON state at signal concentrations above that threshold. Any input states which are below that threshold will be OFF and any above will be ON. The bandpass produces three partitioned regions rather than two; OFF at low signal concentrations, ON at medium concentrations, and OFF again at high concentrations. The lowpass and bandstop are inversion of the highpass and bandpass.

There are some constraints on how the state space can be partitioned by the activation functions. The 00 state must be OFF for the highpass and bandpass, otherwise these would be constitutively on and lowpass functions respectively. Similarly the 00 state must be on for the lowpass and bandstop, otherwise these would be constitutively off and highpass respectively. However, there is no comparable constraint for the 11 state. For example, the highpass can be OFF in the 11 state if the two signal sources are too far away to produce a signal over the concentration threshold.

#### 1.3 All two-input logic gates can be realised with a single receiver

Once we know the allowable orders of the input states and how our activation functions partition that space, we can demonstrate how this paradigm can produce two-input logic gates. First we take the highpass function (OFF|ON). For each admissible position of the partition boundary we will enumerate all the digital functions that can be obtained by swapping the order of input states. There are five possible positions of the partition boundary we need to consider. $|00, 10, 01, 11$ is not possible for the reasons stated above. For the $00|01, 10, 11$ position we can see that we obtain an OR logic gate - output is on with either or both inputs are on. If we swap the order of 10 and 01 - $00|10, 01, 11$- we still produce an OR logic function as the same input states are above the ON threshold. For the next boundary position; $00, 01|10, 11$ is distinct from $00, 10|01, 11$, so we gain two logic gates - A and B. Similarly to the OR gate, $00, 01, 10|11$ and $00, 10, 01|11$ both produce and AND gate, and $00, 01, 10, 11|$ and $00, 10, 01, 11|$ both produce and OFF gate. This means that with the highpass we have a total of five logic gates: OR, A, B, AND, OFF (Supplementary Figure 1A). With the lowpass we have the same boundary positions, but with the inverse mapping (ON|OFF) and through similar arguments, or through inversion of the five highpass logic gates, we gain another five gates (NAND, NOT A, NOT B, NOR, ON) (Supplementary Figure 1B). This means that with the highpass and lowpass we can achieve 10 of the 16 two input logic gates.

In order to achieve all sixteen two-input logic gates, we require the bandpass and bandstop as they are able to isolate states at intermediate concentrations. Without this capability, we could not perform IMPLY or XOR logic. Interestingly, we could perform XNOR and IMPLY logic using the methods described later for multiple receivers. You can also note that the bandpass and bandstop replicate all of the logic gates that can be encoded by the highpass and lowpass respectively. We retain the highpass and lowpass in our set of activation functions as they are easier to engineer and may prove more robust for certain functions.

#### 1.4 Generation of valid orders of input states

For two-input logic, there are $4! = 24$ ways of ordering the input state, but only two of those orderings meet our constraints: $00, 01, 10, 11$ and $00, 01, 10, 11$. For three-input logic these numbers explode; there are now eight input states rather than four (000, 001, 010, 011, 100, 101, 110, 111), leading $8! = 40320$ possible orderings. We have similar constraints to the two-input case:

$$
\begin{aligned}
A_{000} &\leq A_I, A_{II}, A_{111} \\
A_{001} &\leq A_{101}, A_{011} \\
A_{010} &\leq A_{011}, A_{110} \\
A_{100} &\leq A_{101}, A_{110} \\
A_{000}, A_I, A_{II} &\leq A_{111}
\end{aligned}
$$

where the set of states with one and two inputs ON will be labelled as $I = \{001, 010, 100\}$ and $II = \{011, 101, 110\}$ respectively. Furthermore, in the three input case there are additional higher order constraints, for example if group $I$ is ordered such that,

$$A_{100} < A_{010} < A_{001}$$

this imposes a constraint on group $II$,

$$A_{110} < A_{101} < A_{011}$$

##### 1.4.1 Algorithmic generation of valid orders

We developed a method to generate all the allowable orders of input states for a given number of inputs. Here an input state is represented by a string of letters, so that we don't have to type out long

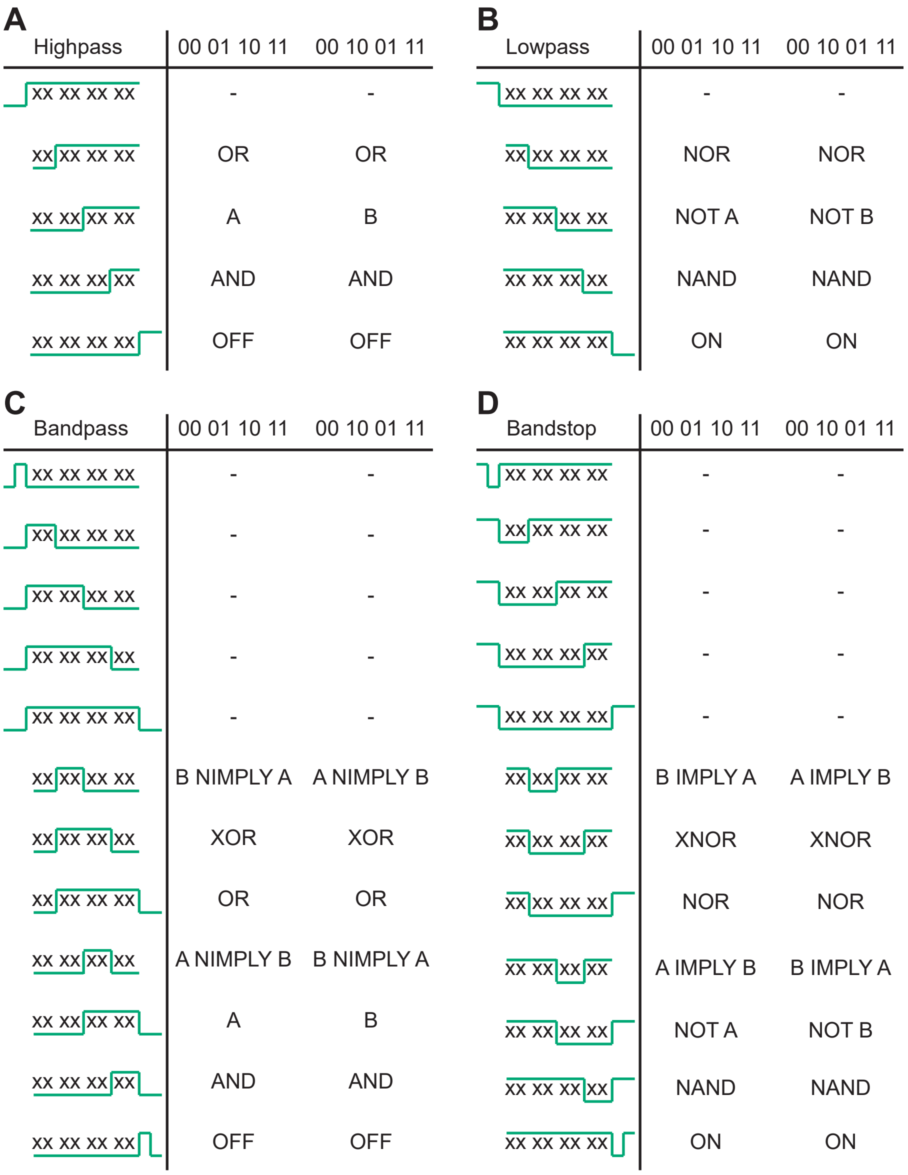

Figure 1: Activation function partitioning of input state space for two-input logic. There are two allowable orderings of the input states in order to maintain the constraint that 00 is lowest and 11 is highest. For each of these orders, we can partition the states into ON and OFF using one of the four activation functions.(A) Highpass can produce five of the sixteen two-input logic functions. (B) Lowpass produces the inverse of the highpass functions. (C) Bandpass and (D) bandstop allow us to produce all sixteen two-input logic functions.

strings of 1s and 0s, where each letter represents an input that is turned on. All inputs included in the string are on and those not included in the string are turned off. For example ADE could be an input state for a five-input gate where input A,D and E are turned on and inputs B and C are turned off. The method generates all possible sequences of input states from lowest to highest signal concentration. The algorithm uses the existing information from input states previously assigned to a given sequence to generate the subsequent states. This ensures that no invalid sequences are generated.

The methods is split into two stages, the first stage is iterative and works as follows. At each step the function considers an incomplete sequence of input states and generates all valid options for the input state at the next position. The valid options for the next state are found by combining states already in the sequence and for each possible combination we record all indices of the input states which can produce that combination. This works as follows (Algorithm S1)

1. The first non-zero state in the current sequence (first state) is combined with the item after it (second state) by concatenating both input states and sorting alphabetically. If the combined state already exists in the current sequence, increment the index of the second state by one so that the next position in the sequence is considered as the second state. For any combined state not already in the sequence we store the index of the first state and second state, if this combined state is found again by subsequent combinations we also store these indices.

2. We then increment the pointer to the first state by one and repeat until there are no more states to be combined and return the potential next states and the indices which were combined to produce them.

**Algorithm 1** get_next_states

```
input: current_sequence
poss_next_states =
for i, s1 in enumerate(current_sequence) do
    for j, s2 in enumerate(current_sequence[i:]) do
        if set(s1) ⋂ set(s2) == None then    ▷ if no inputs will be repeated in the combined state
            next_possible_state = combine_and_sort(s1 ⋃ s2)  ▷ Combine the two states and sort in alphabetical order
            if next_possible_state not in current_sequence then
                pos_next_states[next_possible_state].append([i,j])    ▷ Add the indices of the combination
return poss_next_states
```

The second part of the function is to apply constraints to the possible states generated in the previous algorithm (Algorithm S2). For some sequences the next possible states will be constrained by the existing sequence. For example, for the sequence 0, B, D, A, BD, C, AB, AD, two of the possible next states are BC and ABD. BC can be generated from combining the states at indices 1,5 (B and C) and ABD from states at 3, 4 (A and BD) and 2, 6 (D and AB). Now, just looking at BC from (1,5) and ABD at (3,4) we cannot exclude either state. However if we also consider the second pairing of (2,6) which also produces ABD we see that ABD cannot be the next in the sequence, because D > B and AB > C then ABD > BC and ABD cannot go next.

**Algorithm 2** apply_constraints

```
input: possible_next_states
states = possible_next_states.keys()
valid_states = []
for each combination of: state1, state2 in states do
    if ind1[0] ≤ ind2[0] and ind1[1]≤ ind2[1] for each combination of indices ind1, ind2 in possible_next_states[state1], possible_next_states[state2] then    ▷ state1 is a valid next state
    valid_states.append(state1)
return valid_states
```

The final algorithm (Algorithm 3) combines both the previous algorithms. Starting from a list of the single input states, for a given number of inputs it will repeatedly apply Algorithms 1 and 2 to elements of a queue containing all possible orders until no more operations can be completed and all possible orders of states are generated.

**Algorithm 3** generate_valid_orders

```
input: single_input_list
global queue                                          ▷ queue_order contains all the list of orders
queue.put([])                                         ▷ start with empty sequence
while True do
    sequence = queue.get()                            ▷ Get the first list of order in the queue
    poss_next_state = get_next_states(sequence)
    valid_next_states = apply_constraints(poss_next_states)
    single_next_states = single_input_list - set(sequence)    ▷ Single input states haven't appeared can be the next possible state
    all_next_state = single_next_states ⋃ valid_next_states   ▷ All the next possible states
    if all_next_state is not Empty then
        for item in all_next_state do
            queue.put(list_of_order+[item])           ▷ Add one of the next possible state to the end of the list, and put the list to the end of the queue_order
    if all_next_state is Empty then
        return queue                                  ▷ all possible orders of states
```

Using the Algorithm 3, we can determine the valid input state orderings for a given number of inputs (Supplementary Table 1)

Table 1: Number of possible input state orders that meet the constraints of our paradigm

| Number of inputs | Number of possible ordering |
|---|---|
| 1 | 1 |
| 2 | 2 |
| 3 | 12 |
| 4 | 336 |
| 5 | 65520 |

#### 1.5 One receiver cannot realise all three-input digital functions

After the generation of orders, we can then apply the activation functions to get all the logic gates for 1 output. We apply every possible bandpass and bandstop function to the all generated sequences to find the logic functions which can be constructed with one receiver. There are 12 valid orders of input states for three-input logic. For the single partition activation functions there are 8 positions at which we can partition the state space. This results in 96 order-partition combinations, but only 19 of those result in unique logic (because, for example, $00, |01, 10, 11$ is equivalent to $00, |10, 01, 11$). For the bandpass and bandstop there are 29 ways to position the state space, resulting in 348 order-partition combinations, of which 76 result in unique logic. From this, it is clear that we cannot produce all 256 possible three-input logic functions if we rely on a single receiver colony to produce an output. Supplementary Table 2 contains the number of possible logic gates that can be generated with 1 to 5 inputs.

Table 2: Number of possible logic gates with a single receiver.

| Number of inputs | All gates = $2^{2^n}$ | Achievable gates |
|---|---|---|
| 1 | 4 | 4 |
| 2 | 16 | 16 |
| 3 | 256 | 152 |
| 4 | 65536 | 4034 |
| 5 | 4294967296 | 347752 |

#### 1.6 OR operation for multi-receiver logic

Due to the constraints of signal concentration, it is impossible to produce all logic gates with 1 receiver for more than 2 inputs. We investigate whether we can apply logic optimisation to produce complex logic from the OR of simpler logic functions. To obtain the possible two receiver logic gates an OR operation was performed on each single receiver gate with every other single receiver gate. To obtain the three receiver gates each OR operation was done between each single receiver gate and each two receiver gate and so on. The results are shown in Supplementary Table 3.

Table 3: Number of logic gates from each number of outputs

| Number of inputs | 1 receiver | 2 receivers | 3 receivers | Total |
|---|---|---|---|---|
| 1 | 2 | - | - | 2 |
| 2 | 16 | - | - | 16 |
| 3 | 152 | 256 | - | 256 |
| 4 | 4034 | 56416 | 65536 | 65536 |

##### 1.6.1 Conjecture of the upper bound of number of output colonies

When the number of inputs increases to 5, the running time becomes too long and the memory consumption becomes too large, therefore the computation can only do up to 5 inputs 2 outputs case.

Based on the results in Table S3 we conjecture that the maximum number of receivers required for an $n$ input function is $n - 1$, where the one-input case is a trivial corner case. However, we have yet to prove this.

The conjectured relative complexity of our approach was assessed by comparison with two other implementations of spatially distributed biocomputers [1, 2]. The first approach is based on the separation of cells into a set of connected growth chambers [1]. The maximum number of independent growth chambers, named modules, was used for the comparison. A subsequent paper implemented similar circuits on branched 2D circuits on pieces of paper [2] and the maximum number of branches was used for comparison. Both of these quantities scale by $2^{n-1}$. It is shown in Supplementary Figure 2 that as the number of inputs increases the complexity of our approach scales favourably. This means that the distributed circuits considered here are capable of encoding complex digital functions with less biological complexity than the current state of the art.

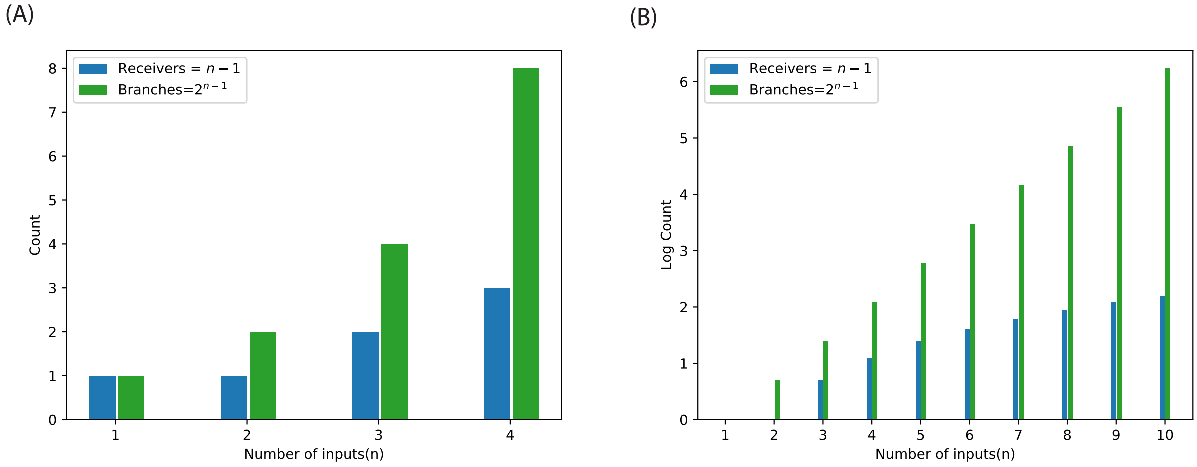

Figure 2: The conjecture result scales for larger number of inputs

#### 1.7 The Macchiato algorithm: optimally distributed spatial circuits

The output of the Macchiato algorithm can then be used to calculate the required pattern of inputs and receivers using a calibrated finite difference model. Designing a spatial digital function, given a truth table, works as follows, where step 3 comprises the Macchiato algorithm itself (Figure 3C):

1. The truth table corresponding to the desired digital function is supplied as input to the Macchiato algorithm

2. The Macchiato algorithm: the following steps are repeated until all 1s in the output of the truth table have been assigned

   The output mapping is reordered to maximise a given block. The block to be maximised depends on the priority set by the user. For example, it is often favourable to maximise the number of input states that are mapped to ON by the highpass, in this situation the algorithm will first try to maximise the size of the upper rightmost block.

   The maximised block is assigned to the don't care (DC) set, meaning that subsequent receivers can map them to either ON or OFF.

3. The blocks are assigned to receivers. The mapping of input states to output states by the receivers is the output of the Macchiato algorithm.

4. A spatial configuration of inputs and receivers that results in the truth table is determined using the calibrated finite difference model. If any of the green receivers are activated the output of the function is ON

##### 1.7.1 The relative importance of the activation functions for the Macchiato algorithm

The output of the Macchiato algorithm is an OR of the outputs of a set of colonies. This means it is important that input states that are required to be mapped to OFF are not mapped to ON by any of the output colonies. However, an input state that is required to be mapped to ON can be mapped to OFF by multiple output colonies, and as long as it is covered by another receiver this won't affect the functions output. The highpass and lowpass can only be used if a block of input states that need to be mapped to ON contain ALL 1 and or ALL 0 respectively i.e. the minimum and maximum signal concentration. As a non-zero amount of signal is required to activate the bandpass, the lowpass is required to map the state ALL 0 to ON, whereas the input state ALL 1 could be covered by the bandpass if required. The bandstop is only required where two blocks of input states contain ALL 0 and ALL 1 and is effectively a convenience that will simplify a number of functions. This means that the activation functions are ranked bandpass, lowpass, highpass, bandstop in order of most important to least important. This intuition was demonstrated and the capability of our approach to build the three-input logic gates was analysed. The Macchiato algorithm was applied to each three-input gate using four different sets of available activation functions. For each set of activation functions, the number of bacterial colonies required to build each gate was found (Table 1). With access to all four activation functions the Macchiato algorithm can achieve all three-input logic gates using one or two output colonies. If we remove the bandstop from the available activation functions we retain the capability to do all the three-input logic gates, however a larger proportion now require two output colonies as expected. If we remove the bandpass activation we are unable to construct the majority of the three-input logic gates, demonstrating the importance of the bandpass. This shows that experimental effort should be prioritised to developing first the bandpass and lowpass.

### 2 Experimental methods

#### 2.1 Strains and plasmids

Overnight cultures were grown in 15 mL Falcon tubes with 3 mL M9 glycerol media [1x M9 salts (BD Difco 248510), 2 mM $MgSO_4$, 0.1 mM $CaCl_2$, 0.4 % glycerol, 0.2 % casamino acids (MP Biomedicals 113060012), 0.035 mg $mL^{-1}$ thiamine] with the relevant antibiotics at 37 °C and shaking at 200 rpm.

Table 4: Strains and plasmids used in this work

| Name | Description | Source |
|---|---|---|
| NEB5$\alpha$ | cloning *E. coli* strain | New England Biolabs |
| BW25113 | Keio collection parent strain | [3] |
| sAJM.1506 | Marionette MG1655 | [4] |
| DH10b-Ptac-T7RNAP | *E. coli* DH10b with IPTG inducible T7RNAP on the chromosome | [5] |
| pRG-PhlF | p15A plasmid, ampicillin resistance, constitutively expressing LacI, and PhlF expressed from a Ptac promoter | [5] |
| pPT-T7wt-O4-PhlF | pSC101 plasmid, chloramphenicol resistance, and sfGFP expressed from a PhlF repressible T7 promoter | [5] |
| pQO34 | pBR322 plasmid, ampicillin resistance, LuxI expressed from an araBAD promoter | this work |
| pQO41 | p15A plasmid, ampicillin resistance, sfGFP expressed from a LuxR-AHL inducible promoter (p59m) | this work |
| pLD10 | ColE1 plasmid, ampicillin resistance, constitutive LldR, LuxI expressed from an lldPRD promoter | this work |
| pKW01 | ColE1 plasmid, ampicillin resistance, constitutive LuxR, sfGFP expressed from LuxR-AHL repressible promoter (J107101) | this work |
| IPTG Bandpass | DH10b-Ptac-T7RNAP:[pRG-PhlF + pPT-T7wt-O4-PhlF] | |
| IPTG Highpass | DH10b-Ptac-T7RNAP:[pPT-T7wt-O4-PhlF] | |
| AHL Highpass | sAJM.1506:[pQO41] | |
| AHL Lowpass | BW25113:[pKW01] | |
| Arabinose Sender | BW25113:[pQO34] | |
| Lactate Sender | BW25113:[pLD10] | |

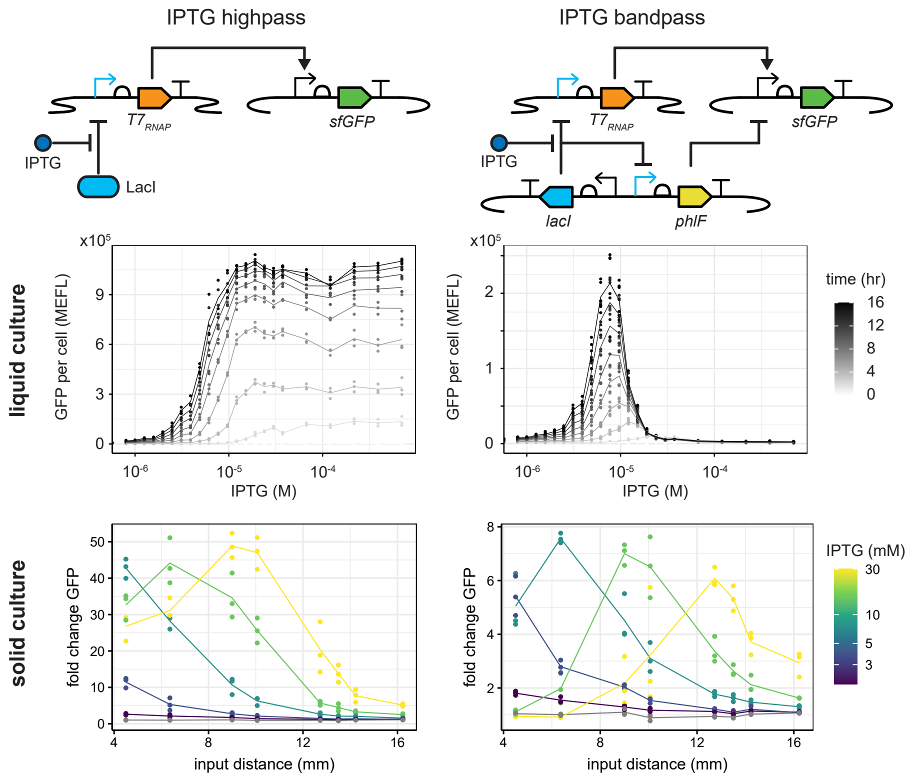

Figure 3: Dose response curves of IPTG inducible highpass and bandpass circuits in liquid and solid culture. Solid culture data is shown at 16 hours.

#### 2.2 3OC6-HSL strain development

We produced a number of strains capable of producing 3OC6-HSL in response to a molecule inducer, or responding to 3OC6-HSL with GFP expression. Most of these strains were produced using golden gate assembly and parts from the CIDAR MoClo [6], from the CIDAR MoClo extension (Richard Murray, Addgene), or the BioBrick library.

#### 2.3 Liquid culture characterisation

The optical density of overnight cultures were measured at 600 nm and the cultures were diluted to an OD600 of 0.05. 120 µL of the diluted culture was pipetted into a 96-well microtitre plate (Greiner 655096). The plate was covered with a plastic lid and incubated in the plate reader (Tecan Spark) for 2 h at 37 °C and shaking. The plate was removed from the plate reader after 2 h and IPTG was added to the wells using a liquid handling robot (Dispendix iDot). The volume of each well was normalised to 125 µL with M9 media. The plate was then sealed with a breathable membrane (Breathe-Easy Z380059) and placed back into the plate reader for a further 16 h at 37 °C and shaking. Measurements of absorbance (600 nm and 700 nm) and fluorescence (excitation 488 nm and emission 530 nm) were taken every 20 min. The data was processed using FlopR [7] to normalise and convert to standard units. The processed data was plotted using custom scripts written in R [8] using packages: ggplot2 [9].

#### 2.4 Agar plate field experiments

5 µL of overnight cultures were diluted in 5 mL of fresh M9 media with the relevant antibiotics. The cultures were then allowed to grow for 2 h at 37 °C and shaking at 200 rpm. 1x M9 agar was made by mixing equal volumes of warmed 2x M9 media and molten 3% agar, and kept in a water bath at 50 °C

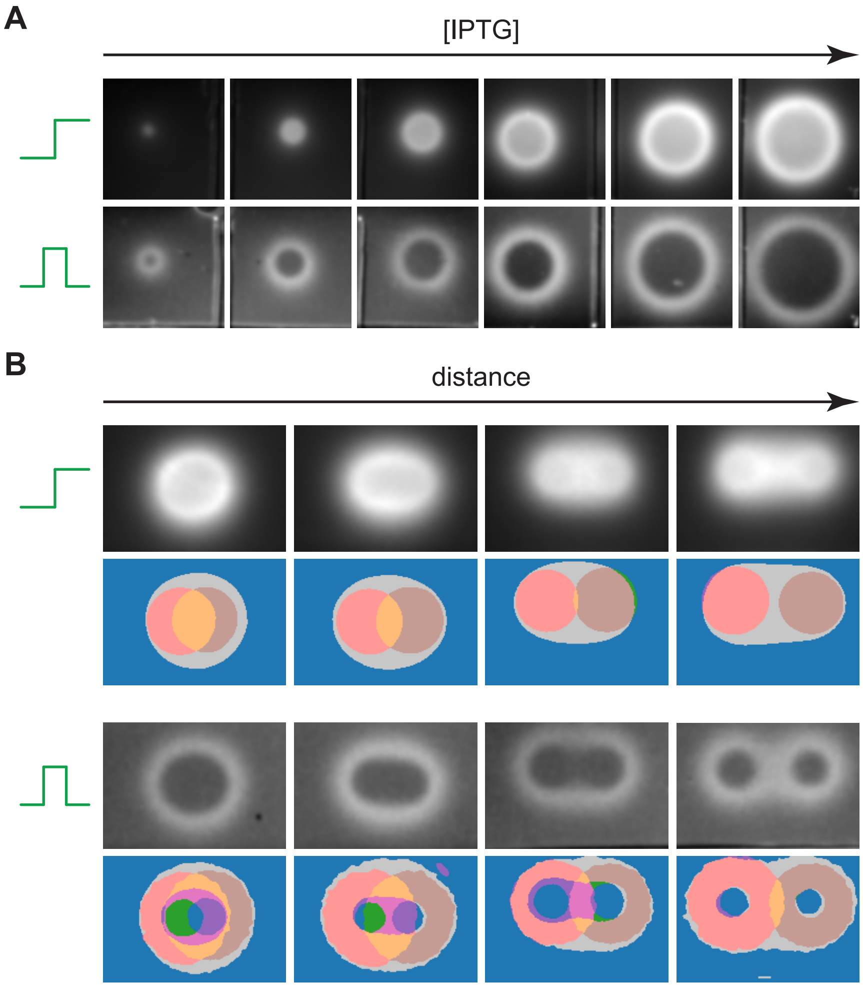

Figure 4: Field experiments exploring the effects of IPTG concentration and distance between inputs. A) Decreasing IPTG concentration from right to left starting at 60 mM with a two-fold reduction each step. B) Increasing the distance between two droplets of IPTG, starting at a distance of 4.5 mm and increasing by 4.5 mm each step. Images shown here were processed as described in the Experimental Methods section. Calculated logic functions at each position are shown below.

to prevent it solidifying. 20 mL of the M9 agar was dispensed into a onewell plate (Greiner 670102) and allowed to dry for 30 min. A 10 mL layer of M9 agar, inoculated with the relevant strain, was then poured over the top. This layer consisted of 5 mL of 2x M9 media with antibiotics, 2.5 mL of 3% agar, and 2.5 mL of culture. This results in an agar density of 0.75 % which makes it easier to produce a level layer before it solidifies and enables the bacterial lawn to grow more evenly. The plates were left to dry for a further 15 min. Channels were cut in the agar to split the plate into four, equally sized, separated areas; roughly between columns 6 and 7, and between rows D and E of a 96-well plate layout. This was to prevent diffusion of inducer between each region of the plate, enabling four experiments to be run concurrently. 1 µL droplets of 7.5 mM IPTG were dispensed at precise locations onto the surface of the bacterial lawns using a liquid handling robot (Opentrons OT2). The plates were left to dry until the droplets were no longer visible; approximately 15 min. They were then placed into an incubator at 37 °C and imaged with a custom imager from loopbio at 0, 16, 18 and 20 hours for growth (red light, intensity 0.7, exposure 4000) and fluorescence (blue light, intensity 3, exposure 20000).

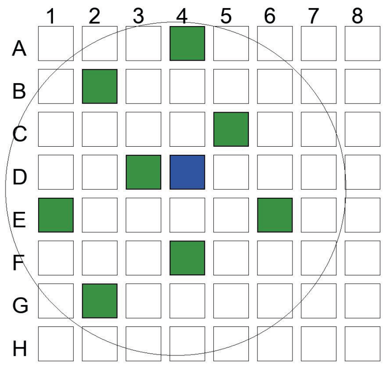

Figure 5: The layout of the characterisation experiment. The inducer is pipetted in the centre of the well (blue) and eight receivers are pipetted at different distances (green)

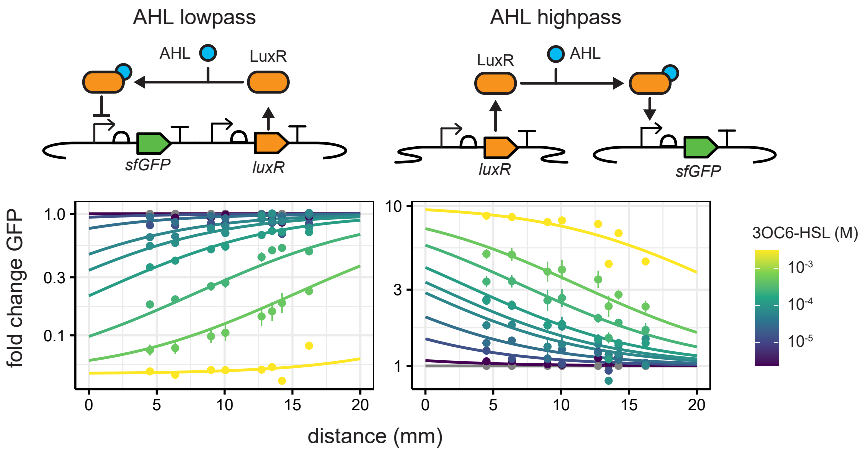

Figure 6: Characterisation of the 3OC6-HSL sensitive receiver strains. A 1 µL droplet of 3OC6-HSL at the given concentration is dispensed as the input. Points and error bars show the mean and standard error of three replicates at 20 hours. Lines show hierarchically fitted Hill functions.

#### 2.5 Agar plate colony experiments

The wells of a 6-well plate (Greiner 657185) were filled with 3 mL of 1x M9 agar made as described above. The plate was allowed to dry for 20 min. The optical density of overnight cultures was measured at 600 nm and the cultures were diluted to an OD600 of 0.3. For the characterisation experiments,1 µL of the diluted culture was dispensed, at the positions shown in Supplementary Figure 5, onto the surface of the agar using a liquid handling robot (Opentrons OT2). For the logic experiments, dispensing positions were determined using the Macchiato algorithm described above. 1 µL of IPTG, at 7.5 mM for logic experiments and at various concentrations for characterisation experiments, was dispensed in the same way. For the characterisation experiments, the plates were left to dry until the droplets were no longer visible; approximately 15 min. For the logic experiments, some IPTG locations were on top of the culture locations. As such, we first dispensed the culture and allowed it to dry until the droplets were no longer visible. Then we dispensed the IPTG and again allowed the plate to dry until the droplets were no longer visible. They were then placed into an incubator at 37 °C and imaged with a custom imager from loopbio at 0, 16, 18 and 20 hours for growth and fluorescence as described above.

#### 2.6 A note on Opentrons dispensing

We noticed greater variation in replicate experiments than we expected from biological noise alone. After extensive investigation, we discovered that the Opentrons was not dispensing volume consistently

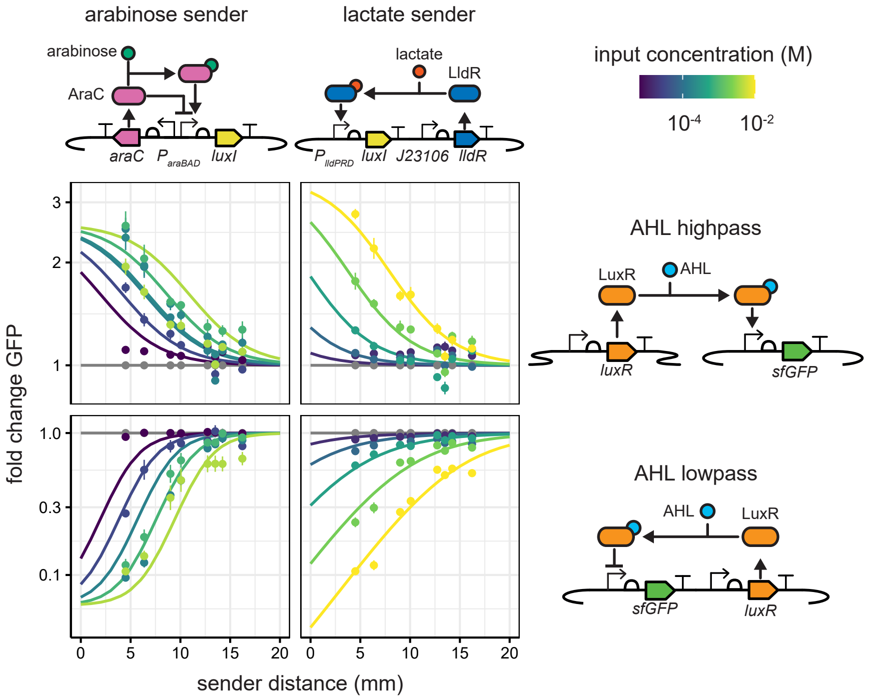

Figure 7: Characterisation of the 3OC6-HSL producing sender strains. A 1 µL droplet of sender culture was placed at the centre of each well. The agar was impregnated with the given concentration of the respective inducer. Each combination of sender and receiver was characterised independently i.e. the top left panel shows the arabinose sender with highpass receiver. Points and error bars show the mean and standard error of three replicates at 20 hours. Lines show hierarchically fitted Hill functions. Note that at high arabinose concentrations the model does not fit the data well, as the high concentrations led to growth defects in the sender colonies.

8. On aspirating 10 µL and then sequentially dispensing 1 µL volumes, the first dispensed droplet was consistently larger than the following droplets. Further, the final two droplets were consistently smaller than the prior droplets. As such, we altered our Opentrons protocols to dispense the first droplet to a sacrificial plate, and dispose of the last 2 µL to waste. This small fix considerably reduced experimental variation.

### 3 Image analysis

#### 3.1 Properties of the loopbio imaging platform

The loopbio imaging platform (Supplementary Figure 9) consists of an LED panel containing blue LEDS at 450 nm and red LEDs at 632 nm. These shine light upwards, through a diffuser, to produce

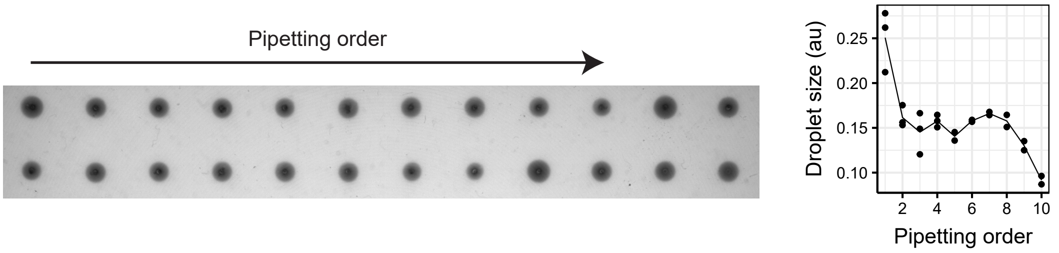

Figure 8: 1 µL droplets of 0.1 % methylene blue. Opentrons aspirated 10 µL and sequentially dispensed until empty. An analogue of droplet size was calculated using our image analysis pipeline, using light intensity as a measure of colony size.

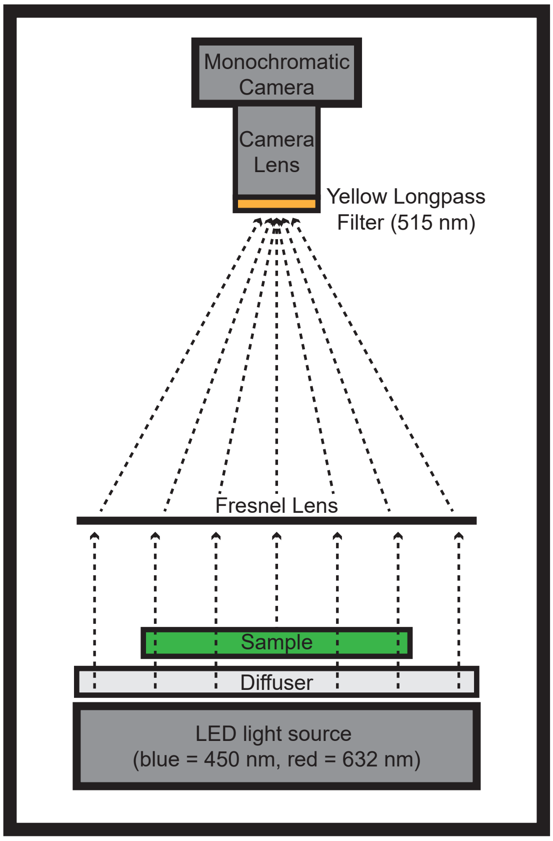

Figure 9: loopbio imaging platform for visualisation of growth and fluorescence of bacterial colonies

an even light source. A sample is mounted in a holder above the diffuser. A Fresnel lens focuses the light from the sample onto the camera lens above. The camera lens is capped with a yellow longpass filter with a cut-on wavelength of 515 nm. This allows the light from the red LEDs to pass through, providing a bright-field image. The light from the blue LEDs is blocked by the yellow filter but fluorescence from the sample induced by the blue light, above 515 nm, will pass through. The brightness of the LED panel can be controlled by adjusting the intensity. The amount of light gathered by the camera can be adjusted by changing the exposure time. The imager is placed in an incubator to maintain the required temperature and remove any background light.

##### 3.2 Field image processing

Fluorescence images from the field experiments were processed with a custom R script. First, a Gaussian blur (standard deviation = 2) was applied to all images. The edges of the images were cropped to remove the rim of the one-well plate. Any gradient was removed from each image by dividing each pixel by its value a 0 h. The images were then normalised by dividing each pixel by the same pixel on an equivalent plate without any inducer. Examples of the resulting images are shown in Supplementary Figure 4. For the distance experiments, the images were then thresholded using the Otsu algorithm. Logic functions were then determined at each position based on the thresholded values from two single input images and a double input image.

##### 3.3 Colony image processing

Extraction of summary pixel information for each region of interest (ROI) on a stack of images is performed by custom software written in R [8] using packages: dplyr [10], imager, scales [11], rlang [12], progressr, future and doFuture [13], foreach [14], stringr [15], tidyr. ROIs are semi-manually

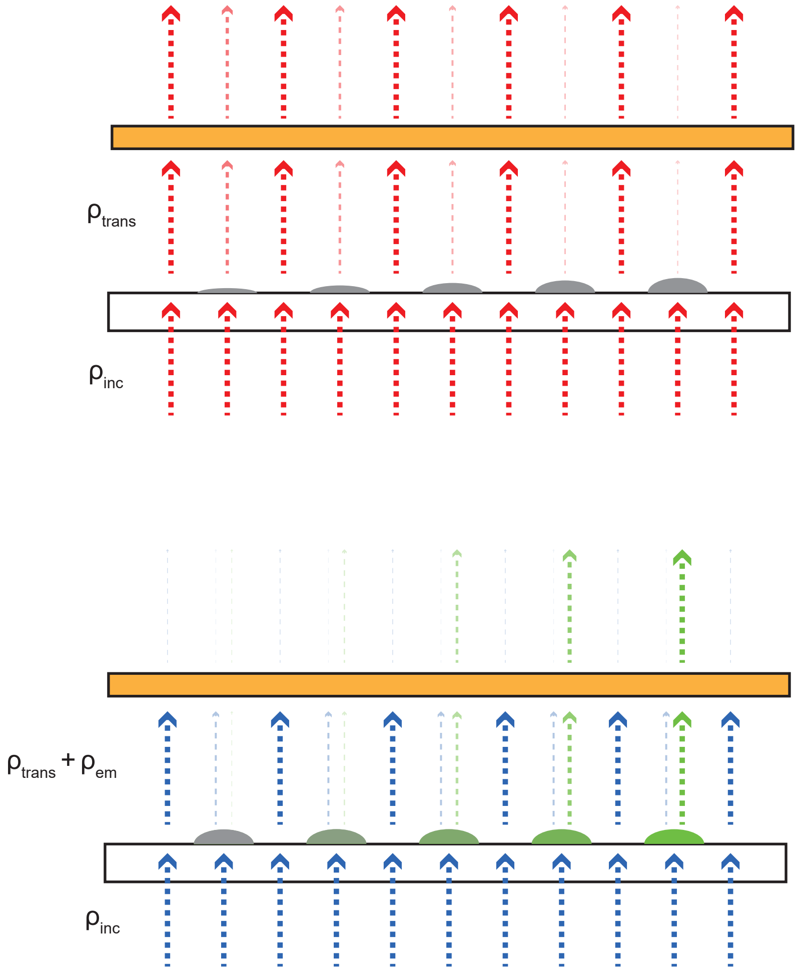

Figure 10: Light transmission through and emission from an illuminated sample.

defined on one representative image in the stack and then applied to all other images. First, a 384-well grid is manually aligned and scaled to a representative image in the image stack. Circular ROIs, centred in each section of the grid, are manually scaled to encompass the bacterial colonies in the representative image. A Gaussian kernel, with standard deviation = 3, is used to blur all images in the stack. The top and bottom 25% of pixels in terms of brightness are discarded. Finally, several statistics are calculated for the remaining pixels within each ROI: mean, median, standard deviation, max, min.

#### 3.4 Calculating growth and GFP values

In the imager, all areas of a plate are imaged concurrently, with different regions of the optics measuring different parts of the plate - each "well" is subjected to a different light source (different part of the LED panel) and is being recorded by a different sensor (different camera pixels) which results in measurements which are position dependent. This means that any parameters that are used to model how measured pixel values correspond to cell count or fluorescence levels, must be calculated or inferred at the pixel (or group of pixel) level rather than for the imager as a whole.

From the pixel values we wish to infer the amount of GFP, $N_{\mathrm{GFP}}$, and the number of cells, $N_{\mathrm{cells}}$ (Supplementary Figure 10). We make the following simplifying assumptions:

- $N_{\mathrm{cells}}(t = 0) \approx 0$

- The light that is incident on the sample $\rho_{\text{inc}} \propto \text{pixel}(t = 0)$

##### 3.4.1 Inferring the number of cells

To calculate the number of cells to an arbitrary unit we use the Beer-Lambert law which states that the concentration of a sample is proportional to the absorbance of the sample. Assuming no scattering by the sample

$$N_{\text{cells}} \propto A = -\log T \quad (1)$$

$$= \log(\frac{\rho_{\text{inc}}}{\rho_{\text{trans}}}) \quad (2)$$

where $A$ is absorbance, $T$ is transmittance, and $\rho_{\text{inc}}$ and $\rho_{\text{trans}}$ are the incident and transmitted light respectively. If we make the same simplifying assumption for transmitted light as we did for incident light i.e. that the measured pixel value is proportional to the light transmitted through the sample

$$N_{\text{cells}} \propto A = \log[\frac{\text{pixel}_{\text{red}}(t = 0)}{\text{pixel}_{\text{red}}(t)}] \quad (3)$$

##### 3.4.2 Inferring GFP concentration

To calculate the concentration of GFP is more complex. Light from the blue LED panel is shone on the sample from underneath, some of that light is attenuated (as with the red light), some of the light excites the GFP in the sample and is emitted at a different wavelength. Any blue light that is transmitted through the sample is mostly, though not completely, blocked by the yellow filter. The green light that is fluoresced is also partially blocked by the yellow filter, though to a much lesser extent. We model the measured light as the sum of the above

$$\text{pixel}_{\text{blue}} \propto \rho_{\text{trans}}\ k_{\text{blue}} + \rho_{\text{em}}\ k_{\text{green}} \quad (4)$$

where the $k$s are coefficients giving the proportion of light of the given colour that can pass through the yellow filter, $\rho_{\text{trans}}$ is the light transmitted through the sample, and $\rho_{\text{em}}$ is the light emitted by the sample. Given our initial assumptions that $N_{\text{cells}}(t = 0) \approx 0$ and $\rho_{\text{inc}} \propto \text{pixel}(t = 0)$

$$\text{pixel}_{\text{blue}}(t = 0) \propto \rho_{\text{trans}}(t = 0)\ k_{\text{blue}} \quad (5)$$

$$\propto \rho_{\text{inc}}\ k_{\text{blue}} \quad (6)$$

We make the simplifying assumption that the absorbance value is the same when using the blue LED as with the red LED. As such

$$A = \log(\frac{\rho_{\text{inc}}}{\rho_{\text{trans}}}) \quad (7)$$

$$\rho_{\text{trans}} = e^{-A}\ \rho_{\text{inc}} \quad (8)$$

$$\propto \frac{\text{pixel}_{\text{blue}}(t = 0)}{e^{A}\ k_{\text{blue}}} \quad (9)$$

We use a model for fluorescence in which

$$\rho_{\text{em}} = \rho_{\text{inc}}\ \phi\ N_{\text{GFP}} \quad (10)$$

where $\phi$ is the detectable emitted light per unit fluorophore per unit incident light and $N_{\text{GFP}}$ is the number of fluorophores. Note here that our $\phi$ encompasses three constants that would normally be included in such a model: fluorescence efficiency, molar absorptivity, and the volume element of detection. Then, from the above equations

$$\text{pixel}_{\text{blue}} = \left(e^{-A} + \frac{k_{\text{green}}}{k_{\text{blue}}}\,N_{\text{GFP}}\,\phi\right)\text{pixel}_{\text{blue}}(t=0) \tag{11}$$

Which gives us $N_{\text{GFP}}$ up to a constant

$$N_{\text{GFP}} \propto \frac{\text{pixel}_{\text{blue}}}{\text{pixel}_{\text{blue}}(t=0)} - \frac{\text{pixel}_{\text{red}}(t)}{\text{pixel}_{\text{red}}(t=0)} \tag{12}$$

### 4 Mathematical modelling

#### 4.1 The finite difference method

The finite difference method solves a system of differential equations by discretising the area of interest and, when using the forward Euler approximation, steps forward through time to simulate dynamics.

The diffusion equation for the concentration of a diffusible molecule $I$ is given by:

$$\frac{\partial I(\mathbf{r},t)}{\partial t} = D\nabla^2 I(\mathbf{r},t) + S(\mathbf{r},t) \tag{13}$$

where $\mathbf{r}$ is the position $t$ is time, $D$ is the diffusion coefficient, $\nabla^2$ is the Laplace operator and $S(\mathbf{r},t)$ is a term that accounts for sources of the molecule. To solve this, the domain of interest is split into a finite number of discrete grid points and an approximate solution is calculated for each point on the grid. The finite difference for a forward time, central space approximation for the diffusion equation in two dimensions is:

$$\frac{I_{i,j,k+1} - I_{i,j,k}}{\Delta t} = D(\frac{I_{i-1,j,k} - 2I_{i,j,k} + I_{i+1,j,k}}{\Delta x^2} + \frac{I_{i,j-1,k} - 2I_{i,j,k} + I_{i,j+1,k}}{\Delta y^2}) + S_{i,j,k}$$

where $I_{i,j,k}$ is the concentration of diffusible molecule at spatial coordinates $i,j$ and timestep $k$, $\Delta x$ and $\Delta y$ are the size of the discrete points in the $x$ and $y$ directions and $\Delta t$ is the timestep between successive iterations. This can be written as:

$$\mathbf{I}_{k+1} = \mathbf{I}_k + \Delta t(H\mathbf{I}_k + \mathbf{S}_k)$$

where $\mathbf{I}_k$ and $\mathbf{S}_k$ are vectors of the current concentration and source terms at all the discretisation points at timestep $k$ respectively and $H$ is a matrix which operates on $\mathbf{I}$ to give the central space difference relations. Using this formula we can start from an initial concentration of diffusible molecule,$\mathbf{I}_0$, and iterate through time. The production of additional molecules by bacterial colonies can be included via the source term $S_k$. This method can be used to model the production of and response to multiple diffusible communication molecules by bacterial colonies.

#### 4.2 Simple finite difference model

A simple finite difference model was developed to verify whether the four activation functions could be used to represent digital logic gates where the inputs are represented by instantaneous sources of diffusible molecules. A square domain was discretised into 200 slices in both the a and y directions to give 40,000 square regions over which the diffusion equation was solved. To simulate the inputs instantaneous sources of diffusible molecule were applied to the initial conditions of the simulation. In this model all units of concentration, distance and diffusion speed were arbitrary. The four activation functions are represented by their digital approximations and assign each square region to OFF or ON as a function of the concentration of signalling molecule in that region. For each of the input states 00, 01, 10 and 11 a simulation was done and each activation functions was applied to the resultant concentration fields. The activation patterns of each activation function were combined to map the resulting logic gates onto each position on the simulation area. This effectively results in a map of the two-dimensional area around the inputs, where the colour of each region represents the logic gate that would be encoded by an receiver if it was placed within that region.

#### 4.3 Calibrated finite difference model

To design logic gates we developed a more complex finite difference model which models cell growth and production of GFP using realistic activation functions.

##### 4.3.1 Zong et al. steady-state model

In the paper, the system is modelled using Hill like functions with some adaptation. The output from each of the two $P_{\mathrm{TAC}}$ promoters is described by

$$Output = \alpha \frac{\left(1 + \left(\frac{Input_I}{K_I}\right)^{n_I}\right)}{1 + \left(\frac{Input_I}{K_I}\right)^{n_I} + K_{lac}} + \beta \tag{14}$$

where $Input_I$ is IPTG and each promoter has different parameters to take account of their differing context. The output from these promoters is T7 (*Activator*) and PhlF (*Repressor*). The output of the third promoter is GFP which is given by

$$GFP = \alpha \frac{\left(\frac{[Activator]}{K_A}\right)^{n_A}}{1 + \left(\frac{[Activator]}{K_A}\right)^{n_A} + \left(\frac{[Repressor]}{K_R}\right)^{n_R}} + \beta + \left(\frac{[Activator]}{K_A}\right) \delta_R \tag{15}$$

##### 4.3.2 Simple dynamic model

We will reuse most of the steady-state model but add a dilution term due to bacterial growth and division. We can even reuse most of the parameters with some simple modifications. Let's assume we have a simple model of gene expression in which $x$ is expressed from an inducible promoter at a maximal rate $k_\alpha$ which is modified by some function $F$ (in our model a Hill function), there is some leaky expression at a rate $k_\beta$, and the cells are growing and dividing at a rate $\mu$. The expression for the change in $x$ over time $t$ is

$$\frac{dx}{dt} = k_\alpha \; F(I) + k_\beta - \mu(t) \; x \tag{16}$$

If we assume that transcription factor binding/unbinding occurs much faster than transcription and translation, we can say that $F(I)$ is only dependent on input $I$ and not on $t$. As such, at steady state

$$0 = k_\alpha \; F + k_\beta - \mu \; x \tag{17}$$

$$x = \frac{k_\alpha}{\mu} \; F + \frac{k_\beta}{\mu} \tag{18}$$

$$x = \alpha \, F + \beta \tag{19}$$

So we can get the dynamic production rates by multiplying the already known $\alpha$ and $\beta$ terms, from the Zong steady state model, by the growth rate $\mu$. We should consider that this relationship is only defined at steady state but we will use it across our timecourse. Moreover, we will use the time varying growth rate as this will naturally give us the highest protein expression rates when growth is fastest and reduce expression as cells move into stationary phase. This leads to our dynamic model

$$\frac{d\mathrm{T7}}{dt} = \frac{\alpha_{T7}\ \mu\ \left(1+\left(\frac{\mathrm{I}}{K_{IT}}\right)^{n_{IT}}\right)}{1+\left(\frac{\mathrm{I}}{K_{IT}}\right)^{n_{IT}}+K_{lacT}} + \beta_{T7}\ \mu - \mu\ \mathrm{T7} \tag{20}$$

$$\frac{d\mathrm{R}}{dt} = \frac{\alpha_{R}\ \mu\ \left(1+\left(\frac{\mathrm{I}}{K_{IR}}\right)^{n_{IR}}\right)}{1+\left(\frac{\mathrm{I}}{K_{IR}}\right)^{n_{IR}}+K_{lacR}} + \beta_{R}\ \mu - \mu\ \mathrm{R} \tag{21}$$

$$\frac{d\mathrm{GFP}}{dt} = \alpha_{GFP}\ \mu\ \frac{\mathrm{T7}^{n_A}}{K_A^{n_A}+\mathrm{T7}^{n_A}}\frac{K_R^{n_R}}{K_R^{n_R}+\mathrm{R}^{n_R}} + \beta_{GFP}\ \mu - \mu\ \mathrm{GFP} \tag{22}$$

When $R = 0$ at all time points this models the highpass response, for $R > 0$ this models a bandpass response.

##### 4.3.3 Parameterising growth

The model presented above describes protein expression within single cells. The population size therefore affects protein expression only through its effect on growth rate. We can infer the growth rate directly from the change in density of our cultures. It is, however, more robust to fit a growth model to this data and calculate growth rates from the parametrised model.

One such growth model is the Gompertz model.

$$N(t) = Ae^{-e^{\frac{\mu_m e}{A}(\lambda - t)+1}}, \tag{23}$$

where $A$ is the maximum population size as $t$ approaches infinity, $\mu_m$ is the maximum specific growth rate and $\lambda$ is the lag time. The derivative of this function can be used to calculate the growth rate at time $t$:

$$\frac{dN}{dt} = \mu = \mu_m e^{\frac{\mu_m}{A}(e\lambda - et) - e^{\frac{\mu_m}{A}(e\lambda - et)+1}+2} \tag{24}$$

##### 4.3.4 Fitting the model

To simulate the response of colonies of bacterial to IPTG inputs the finite difference method was used to solve the equations for $\frac{d\mathrm{T7}}{dt}$, $\frac{d\mathrm{R}}{dt}$, $\frac{d\mathrm{GFP}}{dt}$, $\frac{dA(\mathbf{r},t)}{dt}$ and $\frac{dA(\mathbf{r},t)}{dt}$. To simulate a well in the 6 well plate (a circle with diameter 35mm). A square of width 35mm was modelled, and circular boundary conditions were imposed such that IPTG could not diffuse outside of a circle with diameter 35mm. The square was discretised by splitting it into 1521 (39 by 39) squares each with an area of 0.9mm$^2$. In comparison to the toy model above this is a much coarser grained model. This was done because the parametrisation procedure requires many model simulations and a model of greater size would've been prohibitive in terms of simulation time. The decision to split the area into 39 slices on each axis also means that a distance of 4.5mm (the spacing of the Opentron positions) is almost exactly equal to 5 grid points, meaning we can place inputs and receivers in the correct place despite using a more coarse grained model. The configuration of IPTG and receivers used in the characterisation experiments (Figure S5 and the concentration of $N$, $T7$, $R$, GFP and $A$ was solved for each square using the Runga-Kutta finite difference method by calling the solve˙ivp method from Scipy [16]. The output of any given parametrisation of the model can be compared with characterisation data using the sum square error of the predicted GFP levels with the observed florescence. A particle swarm algorithm [17] was used which explored the space of parameters to minimise the sum square error. This was repeated for both the the highpass and bandpass receivers. For the highpass the following

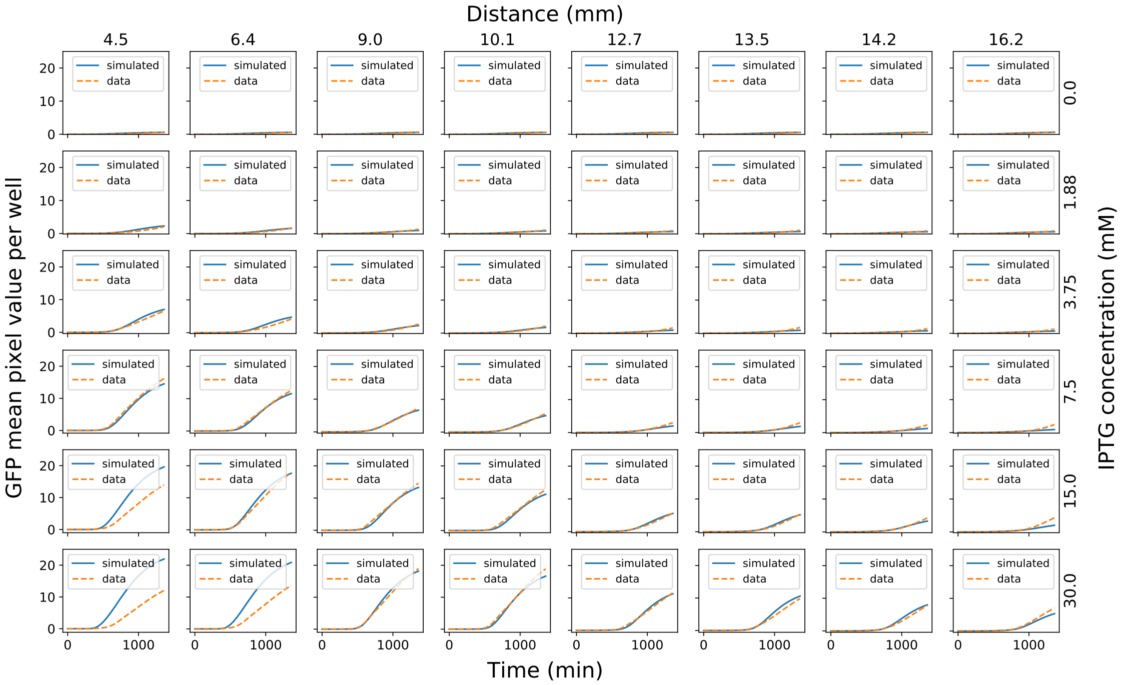

Figure 11: Characterisation data and model fit for the highpass receivers

parameters were fitted: $D_A$, $\alpha_T$, $\beta_T$ $K_{IT}$, $n_{IT}$, $K_{lacT}$, $T7_0$, $\alpha_G$, $\beta_G$, $n_A$, $K_A$, $G_s$. For the bandpass the following parameters were fitted: $D_A$, $\alpha_T$, $\beta_T$ $K_{IT}$, $n_{IT}$, $K_{lacT}$, $T7_0$, $\alpha_G$, $\beta_G$, $n_A$, $K_A$, $G_s$, $\alpha_R$, $\beta_R$, $K_IR$, $n_IR$, $K_{lacR}$, $R_0$, $n_R$, $K_R$.

The results of the model fit for the highpass and bandpass receivers are shown in Figures S11 and S12 respectively. We found that at high concentrations of IPTG the output of the highpass receiver decreases, perhaps due to a toxicity effect. This is not captured in our model and we therefore removed the experiments with (IPTG concentration, distance) combinations of (30mM, 4.5mm) (30mM, 6.4mm) and (15mM, 4.5mm) from consideration when fitting the model. This prevents these combinations from negatively affecting the fit of the model for the other combinations. With exception of these three distance concentration combinations that were explicitly excluded the model shows good agreement with the experimental data for both the highpass and bandpass.

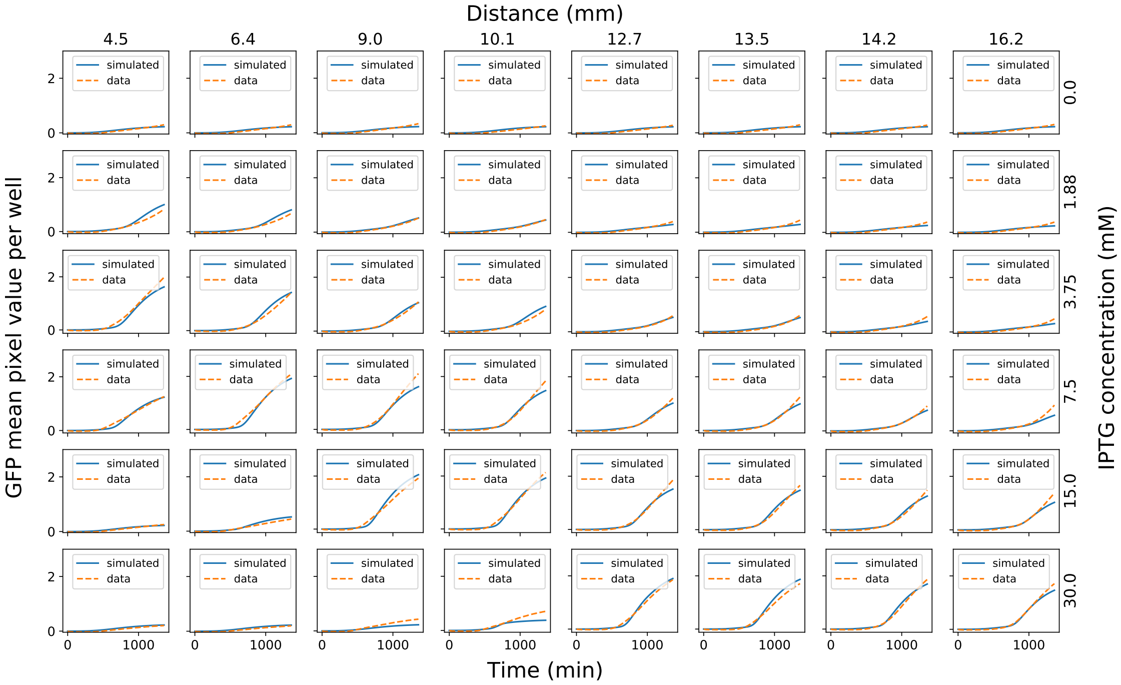

Figure 12: Characterisation data and model fit for the bandpass receiver.

[8] R Core Team. *R: A Language and Environment for Statistical Computing.* R Foundation for Statistical Computing, Vienna, Austria, 2022.

[9] Hadley Wickham. *ggplot2: Elegant Graphics for Data Analysis.* Springer-Verlag New York, 2016.

[10] Hadley Wickham, Romain François, Lionel Henry, and Kirill Müller. *dplyr: A Grammar of Data Manipulation*, 2022. R package version 1.0.10.

[11] Hadley Wickham and Dana Seidel. *scales: Scale Functions for Visualization*, 2022. R package version 1.2.1.

[12] Lionel Henry and Hadley Wickham. *rlang: Functions for Base Types and Core R and 'Tidyverse' Features*, 2022. R package version 1.0.6.

[13] Henrik Bengtsson. A unifying framework for parallel and distributed processing in r using futures. *The R Journal*, 13(2):208–227, 2021.

[14] Microsoft and Steve Weston. *foreach: Provides Foreach Looping Construct*, 2022. R package version 1.5.2.

[15] Hadley Wickham. *stringr: Simple, Consistent Wrappers for Common String Operations*, 2022. R package version 1.4.1.

[16] Pauli Virtanen, Ralf Gommers, Travis E. Oliphant, Matt Haberland, Tyler Reddy, David Cournapeau, Evgeni Burovski, Pearu Peterson, Warren Weckesser, Jonathan Bright, Stéfan J. van der Walt, Matthew Brett, Joshua Wilson, K. Jarrod Millman, Nikolay Mayorov, Andrew R. J. Nelson, Eric Jones, Robert Kern, Eric Larson, C J Carey, İlhan Polat, Yu Feng, Eric W. Moore, Jake VanderPlas, Denis Laxalde, Josef Perktold, Robert Cimrman, Ian Henriksen, E. A. Quintero, Charles R. Harris, Anne M. Archibald, Antônio H. Ribeiro, Fabian Pedregosa, Paul van Mulbregt, and SciPy 1.0 Contributors. SciPy 1.0: Fundamental Algorithms for Scientific Computing in Python. *Nature Methods*, 17:261–272, 2020.

[17] Lester James Miranda. Pyswarms: a research toolkit for particle swarm optimization in python. *Journal of Open Source Software*, 3(21):433, 2018.
